# Leukemia-mutated proteins PHF6 and PHIP form a chromatin complex that represses acute myeloid leukemia stemness

**DOI:** 10.1101/2024.11.29.625909

**Authors:** Aishwarya S. Pawar, Patrick Somers, Aleena Alex, Subin S. George, Charles Antony, Roman Verner, Sanese K. White-Brown, Mohit Khera, María Saraí Mendoza-Figueroa, Kathy Fange Liu, Jennifer J. D. Morrissette, Vikram R. Paralkar

## Abstract

Myeloid leukemias are heterogeneous cancers with diverse mutations, sometimes in genes with unclear roles and unknown functional partners. PHF6 and PHIP are two poorly-understood chromatin-binding proteins recurrently mutated in acute myeloid leukemia (AML). *PHF6* mutations are associated with poorer outcomes, while *PHIP* was recently identified as the most common selective mutation in Black patients in AML. Here, we show that PHF6 is a transcriptional repressor that suppresses a stemness gene network, and that PHF6 missense mutations, classified by current clinical algorithms as variants of unknown significance, produce unstable or non-functional protein. We present multiple lines of evidence converging on a critical mechanistic connection between PHF6 and PHIP. We show that PHIP loss phenocopies PHF6 loss, and that PHF6 requires PHIP to occupy chromatin and exert its downstream transcriptional program. Our work unifies PHF6 and PHIP, two disparate leukemia-mutated proteins, into a common functional complex that suppresses AML stemness.

## Introduction

Myeloid leukemias are a heterogeneous group of hematopoietic cancers with mutations, translocations, or deletions of over one hundred genes or genomic regions^1–3^. Many of these alterations recurrently involve the same protein complexes, like the *RUNX1/CBFB* complex^4^, the cohesin complex (*SMC1A*, *SMC3*, *RAD21*, *STAG1/2*)^5^, the polycomb complex (*EZH2*, *SUZ12*, *ASXL1*)^6^, or components involving the same cellular process, such as splicing (*SF3B1*, *SRSF2*, *U2AF1*, *ZRSR2*)^7^, DNA methylation (*DNMT3A*, *TET2*), or FLT3 signaling (*FLT3*, *PTPN11, NRAS*, *KRAS*)^8^. Such groupings help consolidate diverse mutations into functional clusters, granting insight into convergent mechanisms driving the disease. Nonetheless, like in many cancers, myeloid leukemias also show mutations in isolated genes whose functions and protein partners remain uncertain, preventing them from being assigned to any complex or pathway. Such genes are often placed in generic categories (like “chromatin-binding proteins”) that lack explanatory power, emphasizing the need for studies that might reveal unexpected mechanistic connections between genes once thought to be unrelated.

*PHF6* (Plant Homeodomain Finger protein 6) is an X chromosome gene with recurrent somatic mutations in multiple hematopoietic malignancies, accounting for 3-5% of acute myeloid leukemia (AML) and myelodysplastic syndrome (MDS), 23% of mixed phenotype acute leukemia (MPAL), and 38% of T-cell acute lymphoblastic leukemia (T-ALL) cases^1,2,9–13^. Mouse models of hematopoietic *Phf6* knockout alone show selectively increased hematopoietic stem cell self-renewal without impairment of differentiation, lineage bias, or overt malignancy^14–17^. We have shown that this selective increase is co-opted in a mouse AML model to boost the self-renewal of leukemia-initiating cells (LICs) without affecting their proliferation, cell cycle, or apoptosis; *Phf6* loss instead specifically increases the fraction of LIC progeny that maintain LIC identity after cell division^18^.

Though the cellular phenotypes of PHF6 loss in myeloid and T-lymphoid leukemia mouse models have been characterized in detail^14,15,18–20^, multiple controversies remain about the molecular function or protein partners of this poorly understood protein. PHF6 has nuclear and nucleolar localization signals and is distributed inside the nucleus between the nucleolus and nucleoplasm^21,22^. The protein has two extended PHD (ePHD) domains, each consisting of a zinc finger domain and an imperfect plant homeodomain (PHD)^22^, prompting the reasonable supposition that it acts through interactions with chromatin. However, reported ChIP-Seq studies provide conflicting evidence as to whether it binds to euchromatin or heterochromatin, and whether it activates transcription or represses it^23–26^. PHF6 has also been variably reported to interact with the nucleosome remodeling deacetylase (NuRD) complex^27,28^, DNA damage repair machinery^20,23,29^, and rRNA transcriptional machinery^21,30^. These findings have not always been validated by subsequent reports^31^. Major gaps therefore remain in our understanding of where PHF6 binds, what effects it exerts on bound loci, what functional partners it requires to produce these effects, and whether any of its functional partners are also mutated in leukemia.

In this study, we use *PHF6* knockout and rescue systems in a human AML cell line to show that PHF6 represses a stemness gene program and promotes differentiation. ChIP-Seq reveals high-confidence PHF6 binding to open and active promoters and enhancers, in a pattern matching active histone marks. Both knockout and rescue systems show that PHF6 represses the transcription of bound genes. To answer unresolved clinical questions about somatic PHF6 missense mutations, we perform a comprehensive dissection of nine mutations, eight of which are classified as “variants of unknown significance” (VoUS) by the current clinical workflow at our university hospital. We find that all of them produce a loss of function due to a combination of reduced protein abundance and compromised chromatin occupancy. We also define a critical functional partner for PHF6: an E3 ubiquitin ligase substrate receptor protein named PHIP (Pleckstrin Homology domain Interacting Protein) that recently came to attention in myeloid malignancies as being mutated in 7% of Black patients with AML^32^. This gene had escaped notice in earlier databases because of mutations in only 0.3% of White patients. Guided by multiple lines of evidence, including the fact that rare germline mutations in both *PHF6* and *PHIP* produce overlapping neurodevelopmental disorders, we identify a novel and direct mechanistic link between the two proteins. We show that PHF6 and PHIP control highly similar transcriptional programs, and that the two proteins form a functional complex on chromatin, with PHF6 unable to occupy chromatin or to exert its downstream transcriptional program in the absence of PHIP. Collectively, our work guides appropriate clinical classification of PHF6 missense mutations, and places PHF6 and PHIP, two disparate proteins, within a common chromatin complex that suppresses AML stemness.

## Materials and Methods

### Cell Culture

THP-1 cells were a gift from Martin Carroll at the University of Pennsylvania and were authenticated using STR sequencing and were cultured in RPMI-1640 (Gibco, 11875085) supplemented with 10% FBS (GeminiBio, 100-106). Cell numbers were maintained between 0.2 to 1 million/ml for optimal growth conditions. ER-HOXB8 cells were a gift from Andres Blanco at the University of Pennsylvania and were cultured in RPMI-1640 (Gibco, 11875085) supplemented with 10% FBS (GeminiBio, 100-106), 2% stem cell factor and 20 mM beta-estradiol. HEK-293T cells were cultured in DMEM (Gibco, A4192101) supplemented with 10% FBS.

For doxycycline treatment, a fresh solution of 250 µg/ml doxycycline (Fisher Scientific, AAJ6057914) was made from a 1 mg/ml stock and added to culture media to achieve the desired final concentration. For experiments with multiple time points of doxycycline induction, cells were plated at the same initial time, and doxycycline was added at different intervals to maintain a common collection time for all samples.

### Proliferation assays

Cells were counted and plated in fresh media at a cell density of 200 k/ml. Cell counts were taken every other day (unless otherwise stated) using the BD Accuri C6 system. Cells were preemptively diluted in fresh media to maintain cell density below 1 million/ml. All proliferation assays were performed in triplicates.

### Generation of PHF6 and PHIP knockout, inducible and mutant clonal cell lines

For generating genomic knockouts, CRISPR guide RNAs were designed using Benchling against exon 4 for *PHF6* and exon 6 and 7 for *PHIP*, and ordered as synthetic guide RNAs (sgRNAs) from IDT. 1.2 pM sgRNA was electroporated along with 0.8 pM recombinant Cas9 ribonucleoprotein (Invitrogen, A36498) into 200,000 cells/reaction using the Neon Transfection System (Invitrogen, MPK3000) with 3 pulses of 1700 V and 20 mA according to manufacturer’s protocol. Single-cell clones were sorted and plated 4 days post-electroporation and verified using Sanger sequencing and immunoblotting. To generate missense mutations of *PHF6*, gRNA and HDR templates were designed using Benchling, and electroporated as detailed above.

For doxycycline-inducible lines (Dox-PHF6 and Dx-R274Q), the wild-type human *PHF6* cDNA sequence was cloned from the human THP-1 cell line, and the *R274Q* mutation was engineered by generating a G to A mutation at nucleotide 821 through InFusion cloning (Takara Biology, 638933) according to manufacturer’s protocol. *PHF6* wild-type and mutant coding sequences were separately cloned into the pCW57-MCS1-2A-MCS2 plasmid (Addgene, 80923) at the EcoR1 site. Lentiviral supernatant was generated using VSVG packaging and used to transduce a *PHF6* knockout THP-1 clone. Single cells were sorted based on dsRed expression (indicating integration of construct), and experimental clones were selected by immunoblotting for the absence of leaky PHF6 protein expression at baseline before doxycycline treatment, and expression of protein (wild-type and mutant) at levels similar to parental THP-1 cells after doxycycline treatment. A *PHIP* knockout subclone was generated from the Dox-PHF6 clone using a gRNA targeted to exon 15 of *PHIP*.

### Lentiviral production and transduction

Vector plasmid (4 µg) and packaging plasmids (2 µg gag-pol (*pCAG-kGP1-1R*), 0.67 µg envelope (*pHDM-G*), and 0.67 µg rev/tat (*pCAG-RTR2*)) were mixed with JetPRIME transfection reagent and buffer (Polyplus, 89129-922) and added to HEK-293T cells in a 10 cm plate. Media was replaced with fresh media 24 hrs post-transfection, and viral supernatant was collected after 72 hrs and used to spin-infect THP-1 cells at 960 x *g* for 90 mins at 37° C with 8 µg/ml of polybrene.

### Immunoblotting

1-2 million cells were washed in DPBS and resuspended in freshly prepared RIPA buffer with protease inhibitors - 1 mM PMSF (Cell Signaling Technology, 8553S), 1X PIC (Sigma, P8340) 1X phosphatase inhibitor (Emsco Fisher, 78427), and 2 mg/ml of chymostatin (Millipore, EI6). In our experience, high-dose chymostatin is essential to prevent PHF6 protein cleavage *in vitro* during lysate preparation. Samples were incubated on ice for 30 mins and centrifuged at 17,000 x *g* for 30 mins at 4° C. Supernatant was collected, and protein concentration was determined by BCA assay (Thermo Scientific, 23225). Protein samples were prepared at a 1.25 µg/µl concentration in 4X Laemmli (BioRad, 1610747) and PAGE was performed on NUPAGE 4-12% Bis-Tris polyacrylamide gels (Invitrogen) and wet transferred overnight to PVDF membranes (BioRad, 620177). Membranes were blocked for 1 hr using 5% (w/v) non-fat milk in TBST, incubated overnight in primary antibody, followed by 1 hr incubation in secondary antibody. Membranes were washed 3 times between each incubation. Images were captured using an Odyssey CLx Imaging System (LI-COR Biosciences, NE, USA), and ImageJ software (Fiji) was used for protein quantification.

### Flow Cytometry

1 million cells were counted, washed, and resuspended in 10 µL of antibody cocktail constituted in cell staining buffer (BioLegend, 420201). After 60 mins of incubation, cells were washed in cell staining buffer and resuspended in 300 µL of fresh buffer. UltraComp eBeads compensation beads (Thermo Fisher Scientific, 01-2222-42) were used for compensation controls. Data were acquired on an LSRFortessa flow cytometry machine (BD Biosciences) and analyzed and visualized using FlowJo (BD Biosciences) and Prism (GraphPad Software).

### Immunofluorescence

Live cells were washed and adhered on lysine-treated slides at 37° C for 1 hr. Once adhered, cells were fixed using 4% PFA (Thermo Scientific, AAJ19943K2) and permeabilized using methanol, followed by blocking in PBS with 10% rabbit serum (Fisher Scientific, 16-120-099) and 0.2% Tween 20 (BioRad, 1706531). Slides were incubated overnight with a primary antibody cocktail diluted in blocking serum, followed by 1 hr of secondary antibody incubation in blocking serum. Finally, cells were incubated with 0.5 µg/ml DAPI before the addition of mounting media (Invitrogen, P10144) and sealing with coverslips. Cells were imaged using an STP800 wide-field microscope (Leica), and quantification was performed using ImageJ software (Fiji) for 30-50 cells across 2-3 slides per clone.

### ATAC-Seq and analysis

50k cells were collected, lysed in 50 ul lysis buffer (10 mM Tris-Cl, 10 mM NaCl, 3 mM MgCl_2,_ 0.1% IGEPAL) to collect nuclei, followed by tagmentation performed using Nextera Tn5 Transposase (Illumina Nextera ATAC-Seq kit, 20034197) according to manufacturer’s protocol. Libraries were prepared using dual-end indexing using NEBNext Ultra II DNA Library Prep Kit (New England Biolabs, E7645L) using the manufacturer’s protocol. The size and concentration of each library were determined using Agilent 2200 Tapestation and KAPA library quantification kit (Roche, KK4824) respectively. Libraries from all samples were pooled in equimolar concentrations. The library pool was quantified using a KAPA library quantification kit and sequenced on NextSeq 500/550 (Illumina) to a depth of 40 million reads per sample. Raw reads were trimmed of adapter sequences using Trimmomatic v0.38^33^ and aligned to the human hg19 genome using Bowtie2 v2.5.0^34^. Peaks of transposase-accessible chromatin were called and quantified using MACS2 v2.2.7.1^35^. Differential and consensus peak analyses were performed using the DiffBind v3.12.0 package in bioconductor^36,37^. Bigwig files were generated using bedGraphToBigWig v302.1^38^.

### RNA extraction, qRT-PCR, RNA-Seq

RNA was extracted from 1 million cells using the Qiagen RNeasy Mini Kit (Qiagen, 74106). β-mercaptoethanol was added to Buffer RLT (10 µl/ml of RLT), and DNase digestion was performed on a column as per the manufacturer’s protocol.

For qRT-PCR, RNA purity and yield were determined using a Nanodrop 1000 spectrophotometer. cDNA from 1ug RNA was prepared using iScript cDNA Synthesis Kit (BioRad, 1708891). Real-time PCR was performed using Luna Universal qPCR Master Mix (New England BioLabs, M3003E) on the QuantStudio™ 6 Real-Time PCR system (Applied Biosystems) and the relative expression levels were calculated using the ΔΔCt method with the average Ct of GAPDH and ACTIN used for normalization.

For RNA-Seq, RNA was quantified using RNA Screentape (Agilent, 5067-5576). 1 µg RNA was used for library preparation using NEBNext Ultra RNA Library Prep Kit for Illumina (New England Biolabs, E7530), and libraries were multiplexed using NEBNext Multiplex Oligos for Illumina (New England Biolabs, E7600). Libraries were quantified using D1000 Screentape (Agilent, 5567-5582) and KAPA Library Quantification Kit (Roche, KK4824). Pooled libraries were sequenced (paired-end) to a depth of 30 million reads per sample on a NextSeq 500/550 (Illumina).

### RNA-Seq Analysis

Raw reads were demultiplexed and FASTQ files were generated using Bcl2fastq v2.20.0.422 (Illumina). Reads were trimmed for low quality or adapter sequences using Trimmomatic v0.38^33^, followed by alignment to the human hg19 genome for THP-1 cells or mouse mm10 genome for ER-HOXB8 cells by STAR v2 aligner^39^. Gene-level read counts were generated using the featureCounts tool from Rsubread v1.6.1^40^. Read count normalization and differential gene expression testing were performed using DESeq2 v1.42.0^41^. Gene set enrichment analysis was performed using UC San Diego and Broad Institute GSEA 4.1.0 software^42,43^. Gene set enrichment data are represented using default parameters and a 0.05 p-value cutoff.

### ChIP-qPCR, ChIP-Immunoblotting, and ChIP-Seq

For PHF6 ChIP-Seq, 40 million cells were collected, washed in DPBS, and double-crosslinked with 1.5 mM EGS (MedSupply Partners, GBS-BC09) for 30 mins at RT on a rotating mixer. Cells were pelleted at 500 x *g* for 5 mins, crosslinked with freshly diluted 1% formaldehyde (Pierce PI28908) for 10 mins at RT on a rotating mixer, and quenched using 125 mM glycine at room temperature. Cells were pelleted and washed 3 times in ice-cold DPBS, rapidly freeze-thawed 3 times using liquid nitrogen, and treated with 4 µl per 40 million cells of MNase (New England Biolabs, M0247S) for 7 mins at 37° C. MNase was inactivated with 10 ul of 0.5 M EGTA (pH 8.0), and nuclei were pelleted at 5600 x *g* for 1 min. The supernatant was discarded, and the nuclear pellet was gently resuspended in 600 µl per 40 million cells of lysis buffer (0.1% SDS, 1% Triton X-100, 10 mM Tris-HCl pH 7.4, 1 mM EDTA, 0.1% Na deoxycholate, 300 mM NaCl, protease inhibitors). The suspension was sonicated in a Bioruptor Pico (Diagenode) at 2 cycles for 30 sec ON/OFF, and centrifuged at 17000 x *g* for 15 mins. The supernatant was collected, 5 μg of Drosophila chromatin per 40 million cells (Active Motif, 53083) was spiked into it, and then 50 µl Dynabeads A (Invitrogen, 10002D) were added and the lysate was clarified overnight at 4° C on a rotating mixer. 10 ug of target-specific antibody plus 2 μg of antibody against Drosophila-specific histone variant H2Av (Active Motif, 61686) was bound to fresh Dynabeads A overnight at 4° C on a rotating mixer. Beads from the clarified lysate were magnetically removed and discarded, and antibody bound beads were added to the lysate, followed by overnight incubation at 4° C on a rotating mixer. Following the incubation, beads were magnetically separated, supernatant was discarded, and beads were washed with low salt (150 mM NaCl, 0.1% SDS, 1% Triton X-100, 1 mM EDTA, 50mM Tris-Cl), high salt (500 mM NaCl, 0.1% SDS, 1% Triton X-100, 1 mM EDTA, 50 mM Tris-Cl) and LiCl (150mM LiCl, 0.5% Na deoxycholate, 0.1% SDS, 1% NP40, 1 mM EDTA, 50 mM Tris-Cl) washes, and then incubated with 200 µl of elution buffer at 65° C on a shaker for 30 mins to elute bound chromatin. After elution, beads were magnetically separated and discarded, and the eluate was de-crosslinked by incubating overnight at 65° C, followed by treatment with 2 ul RNase A (Invitrogen, EN0531) at 37° C and 2 ul Proteinase K (Invitrogen, 25530049) at 56° C for 1 hr each. DNA was purified using a PCR purification kit (Qiagen, 28104) and eluted into 50 µl of nuclease-free water. In our experience, double-crosslinking with EGS and formaldehyde in the described sequence, followed by chromatin fragmentation using MNase (with no more than gentle sonication to rupture nuclear membranes) is essential for efficient PHF6 ChIP. For H3K27ac and PHIP ChIP-Seq, 20 million cells were crosslinked with freshly prepared 1% formaldehyde alone.

For ChIP-qPCR, DNA was diluted 10X, and quantitative PCR was performed using Luna Universal qPCR Master Mix (New England BioLabs, M3003E) on QuantStudio™ 6 Real-Time PCR system (Applied Biosystems) and quantified against standard curve for 5% input sample.

For ChIP-immunoblotting, three parts eluate from beads was boiled with one part 4X Laemmli (BioRad, 1610747) and PAGE was performed on NUPAGE 4-12% Bis-Tris polyacrylamide gels (Invitrogen). Proteins were transferred to PVDF membranes and visualized as described above.

For ChIP-Seq, libraries were prepared using NEBNext Ultra II DNA Library Prep Kit for Illumina (New England Biolabs, E7645) and multiplexed using NEBNext Multiplex Oligos for Illumina (New England Biolabs, E7600). Libraries were quantified using D1000 Screentape (Agilent, 5567-5582) and KAPA Library Quantification Kit (Roche, KK4824). Pooled libraries were sequenced (paired-end) to a depth of 20 to 40 million reads per sample on a NextSeq 500/550 (Illumina).

### ChIP-Seq Analysis

Raw reads were demultiplexed and FASTQ files were generated using Bcl2fastq v2.20.0.422 (Illumina). Reads were trimmed for low quality and adapter sequences using Trimmomatic v0.38^33^. Trimmed FASTQ files were first mapped to the *Drosophila melanogaster* (dm3) genome using Bowtie2 v2.5.0^34^. Spike-in normalization was performed using a published method^44^: A normalization factor was first calculated for each FASTQ file to be normalized by dividing the dm3 genome read number for that file by the smallest dm3 genome read number across all files. The total number of trimmed reads for each file was then divided by the normalization factor for that sample to calculate the normalized number of reads to keep. Each FASTQ file was then sampled for its normalized number of reads to keep, and the sampled FASTQs were mapped to the hg19 genome using bowtie2 v2.5.0^34^. Peak calling was performed using MACS2 2.2.7.1^35^. High-confidence PHF6 and PHIP peaks were defined as statistically significant peaks (padj < 0.05) present in at least two technical replicates. Motif analysis was performed using HOMER^45^ for all ATAC peaks that overlapped with PHF6-bound promoters using non-overlapping ATAC peaks as reference. Bigwig files were generated using bedGraphToBigWig v302.1^38^.

### rDNA Mapping

Trimmed reads were mapped using Bowtie2 v2.5.0^34^ to a customized genome assembly for rDNA mapping (Human_hg38-rDNA_genome_v1.0) previously generated by our lab^46^. Bigwig files were generated using bamCoverage from deepTools v3.5.1^47^.

### Data Visualization and Statistical Analysis

Heatmaps were generated using pheatmap v1.0.12 in RStudio v4.3.2. For volcano plots and regression plots, ggplot2 v3.4.4^48^ package in RStudio was used. PCA plots were generated and peak overlapping was performed using PCA plot w ggplot2 (Galaxy Version 3.4.0+galaxy0)^48^. Metagene plots and profiles were generated using SeqPlots 1.4.1^49^. Tracks were visualized on the IGV browser v2.8.0^50^. All bar graphs were prepared and accompanying statistical analyses were performed using GraphPad Prism v10.0.0 for MacOS (GraphPad Software). Figures and diagrams were generated using Adobe Illustrator and BioRender.

### Clinical genomics classification of PHF6 mutations

PHF6 mutations were reviewed through the standard clinical genomics workflow used by the Penn Center for Personalized Diagnostics for classifying mutations or variants in patient reports. The workflow included review of published functional data, PubMed^51^, ClinVar^52^, gnomAD (v4.1)^53^, COSMIC^54,55^, cBioPortal^56–58^, OncoKB^59,60^, and Varsome^61^, to score and characterize variants into three tiers: (a) Tier 1 - FDA approved therapy associated with the variant, (b) Tier 1 or 2 - potential significance of pathogenicity, and (c) Tier 3 - variant of unknown significance (VoUS), with likely benign or benign variants not reported.

### *In silico* pathogenicity classification of PHF6 mutations

Based on algorithms recommended on the ClinVar portal^52^, we selected the top 4 concordant meta-predictors: REVEL^62^, MetaLR^63^, MetaSVM^64^, and Condel^65^ to predict the pathogenicity of PHF6 mutations using the ePHD2 amino acid sequence (residues 205-333) as input. Note: These algorithms could only be applied to ePHD2 mutations due to the availability of a crystal structure only for that domain. REVEL scores ranged from 0 to 1, and a score of ≥ 0.7 was used as a cut-off for pathogenic variants and a score of ≤ 0.25 was used as a cut-off for benign variants. MetaLR scores ranged from 0 to 1, with higher values more likely to be deleterious. For MetaSVM, negative scores were tolerated by the protein structure and positive scores were deleterious. Condel scores ranged from 0 to 1, and a score above 0.522 was considered deleterious. Mutations with 4 pathogenic meta-predictor scores were classified as Pathogenic, 3 as ‘Likely pathogenic’, 2 as ‘Variant of unknown significance (VoUS)’, and mutations with one pathogenic score or lower were classified as Likely benign.

### Data availability

Datasets from sequencing experiments have been deposited on the Gene Expression Omnibus and are available online - RNA-Seq datasets: GSE281475, GSE281624, GSE281625; ATAC-Seq dataset: GSE281626; and ChIP-Seq datasets: GSE281627, GSE281628, and GSE281629.

## Results

### PHF6 suppresses a hematopoietic stemness gene program and promotes differentiation

We used the ProteinPaint^54,66^ portal to visualize the types and frequencies of *PHF6* somatic mutations in adults with myeloid and lymphoid hematological malignancies (**Fig** 1A). Two-thirds of *PHF6* mutations are frameshift and nonsense mutations distributed throughout the gene body, presumed to produce null alleles due to nonsense-mediated decay. The remaining one-third are missense point mutations clustered in the protein’s second ePHD domain, and the consequences of these missense mutations are unknown. To first study the effect of complete PHF6 loss, we used CRISPR/Cas9 to generate *PHF6* knockout (PHF6^KO^) and wild-type (WT) clones of the THP-1 human AML cell line (**Fig** 1B). Consistent with DepMap^67,68^ CRISPR screen findings (by the Broad Institute) that *PHF6* loss had minimal to no effect on the growth of 26 assayed AML cell lines (**Fig** S1A), we observed that PHF6 loss did not affect the growth of THP-1 cells (**Fig** S1B). This is consistent with our prior report^18^ that *Phf6* loss in mouse AML increases stemness without affecting proliferation. RNA-Seq of human THP-1 clones (**Fig** 1C) showed that HSC and progenitor genes^69^ were positively enriched in PHF6^KO^ compared to WT (**Fig** 1D). To determine whether this shift towards a stem-like transcriptome was accompanied by loss of differentiation, we performed flow cytometry for myeloid markers^70–72^ and observed that PHF6^KO^ clones showed reduced expression of multiple myeloid cell surface markers (**Fig** 1E). To determine whether these effects could be recapitulated in an independent mouse cell line, we knocked out *Phf6* in the mouse ER-HoxB8 GMP cell line^73^ (**Fig** S1C). Similar to human cells, we observed no effect on cell growth (**Fig** S1D). RNA-Seq (**Fig** S1E) showed negative enrichment of differentiated granulocyte genes^74^ (**Fig** S1F) and positive enrichment of self-renewal genes^75^ (**Fig** S1G) in *Phf6^KO^* clones compared to *Wt*, demonstrating a similar shift towards stemness and away from differentiation.

**Figure 1:**
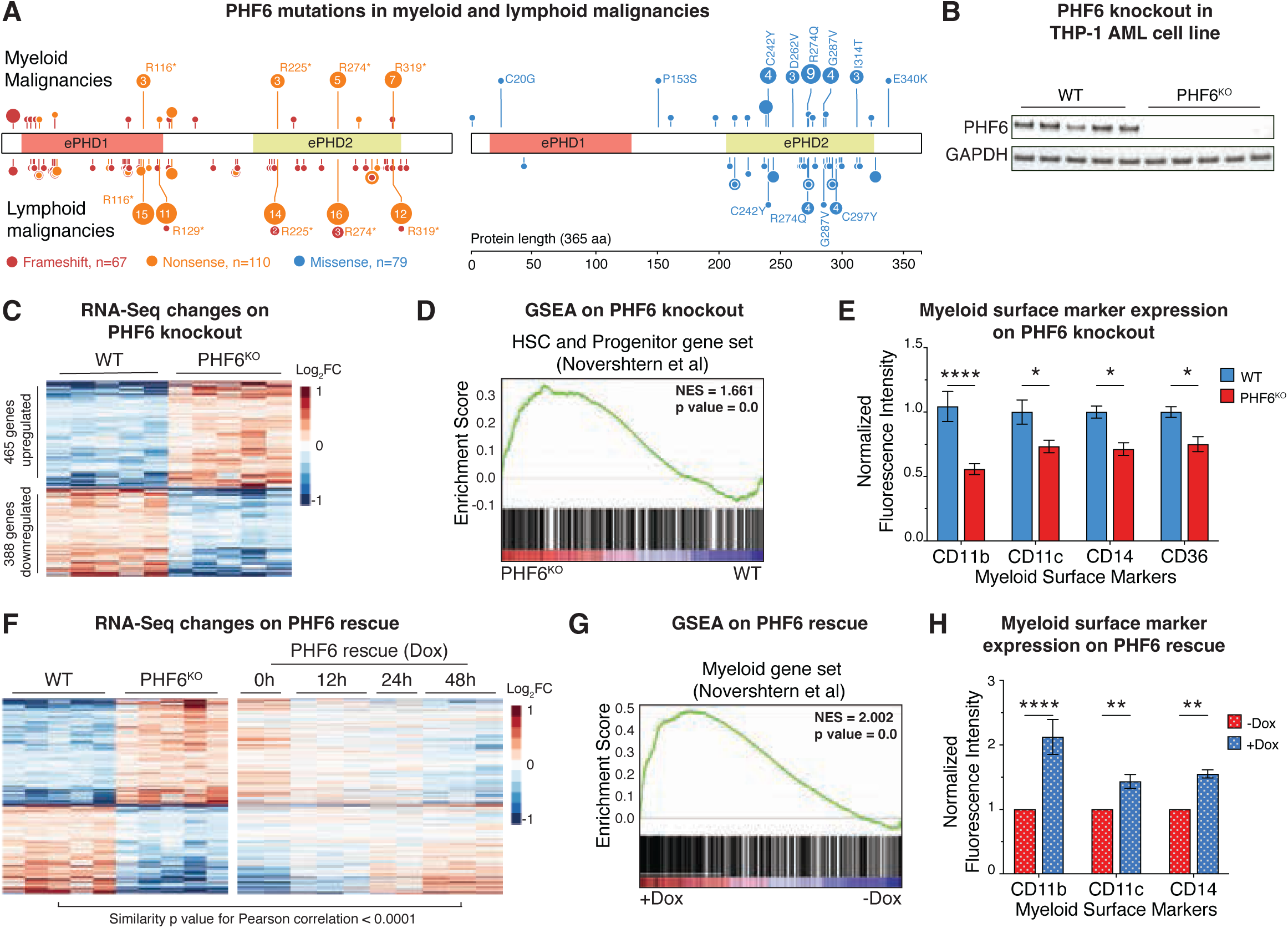
PHF6 suppresses stemness genes and promotes differentiation. A. Lollipop plot of somatic *PHF6* mutations in adults with myeloid (top) and lymphoid (bottom) hematological malignancies. Frameshift and nonsense mutations are shown on the left, and missense mutations are shown on the right. Plot was generated using COSMIC data visualized on the ProteinPaint portal. ePHD1 and ePHD2 domains of PHF6 protein are indicated. B. Immunoblot for PHF6 in WT and PHF6^KO^ THP-1 clones. GAPDH is shown as loading control. C. Heatmap showing 853 differentially expressed genes in PHF6^KO^ compared to WT. D. Gene set enrichment analysis (GSEA) plot showing positive enrichment of HSC and progenitor cell gene set in PHF6^KO^ compared to WT. E. Bar graph showing normalized median fluorescence signal of myeloid surface markers in PHF6^KO^ compared to WT. (n=3) F. Heatmap showing time course of effect of PHF6 rescue on genes differentially expressed in PHF6^KO^ compared to WT. Pearson correlation shows similarity between expression profiles of WT and PHF6 rescue clones at 48 hrs after doxycycline treatment. G. GSEA plot showing positive enrichment of myeloid cell gene set after 48 hours of PHF6 rescue compared to baseline KO state. H. Bar graph showing normalized median fluorescence signal of myeloid markers after 48 hours of PHF6 rescue. (n=3) All bar graphs show mean ± standard error of mean (SEM), ns (not significant) = p ≥ 0.05, *p = 0.01 to 0.05, **p = 0.001 to 0.01, ***p = 0.001 to 0.0001, ****p < 0.0001, by one-way ANOVA with Sidak’s multiple comparison testing.

To determine whether changes produced by PHF6 loss could be reversed on PHF6 re-expression, we developed a doxycycline-driven PHF6 rescue system on the THP-1 *PHF6*^KO^ background (**Fig** S1H). Rescue of PHF6, as expected, showed no effect on cell growth (**Fig** S1I), and RNA-Seq showed progressive reversion over time of the transcriptome to match WT clones (**Fig** 1F, S1J). Relative to the 0 hr time point, the transcriptome at 48 hrs post rescue was positively enriched for myeloid differentiation genes (**Fig** 1G). Surface expression of myeloid markers was also rescued (**Fig** 1H). Collectively, results from both knockout and rescue systems show that PHF6 (likely via a combination of direct and indirect effects) represses stemness genes and promotes differentiation in human AML without affecting proliferation, recapitulating our prior *in vivo* mouse AML findings^18^. This further indicates that THP-1 cells are a suitable human cell system to dissect PHF6 molecular function.

### PHF6 binds gene promoters and represses transcription

PHF6 protein is exclusively nuclear, localizing partly to the nucleolus and partly to the nucleoplasm^21^. The role of the nucleolar fraction is unknown, but nucleoplasmic PHF6 has been reported to bind chromatin, though there is controversy as to whether it binds active or repressed chromatin^23,25^. To resolve this controversy, we sought to define PHF6 peaks by performing multiple replicates of PHF6 ChIP-Seq in the THP-1 line, along with pulldown using the same anti-PHF6 antibody in a PHF6^KO^ clone as a critical negative control in addition to IgG. We identified 8,593 high-confidence PHF6 peaks, almost all of which were at open and active regions of chromatin, evidenced by their alignment with ATAC-Seq and H3K27ac ChIP signal (**Fig** 2A). Overlap of PHF6 peaks with ENCODE-defined cis-regulatory elements (CREs)^76^ (**Fig** S2A) showed that 20% of PHF6 peaks overlapped with loci with promoter-like signatures (PLS), 31% with proximal enhancer-like signatures (pELS, within 2kb of gene promoters), and 36% with distal enhancer-like signatures (dELS) (**Fig** 2B). As per this classification, 51% of PHF6 peaks fell within 2kb of a gene promoter. PHF6 occupancy pattern at bound genes strongly matched that of other active histone marks like H3K4me3, H3K27ac, and H3K9ac, and PHF6-bound genes also showed H3K36me along their bodies, further indicating active transcription (**Fig** 2C). Notably, in contrast with a previous report^23^, we did not observe PHF6 co-localization with repressive marks like H3K27me3 or H3K9me3 (**Fig** 2C). Also in contrast with another report^30^, PHF6 showed no evidence of binding to ribosomal DNA repeats (**Fig** S2B), and hence we did not further investigate any nucleolar role in this work.

**Figure 2:**
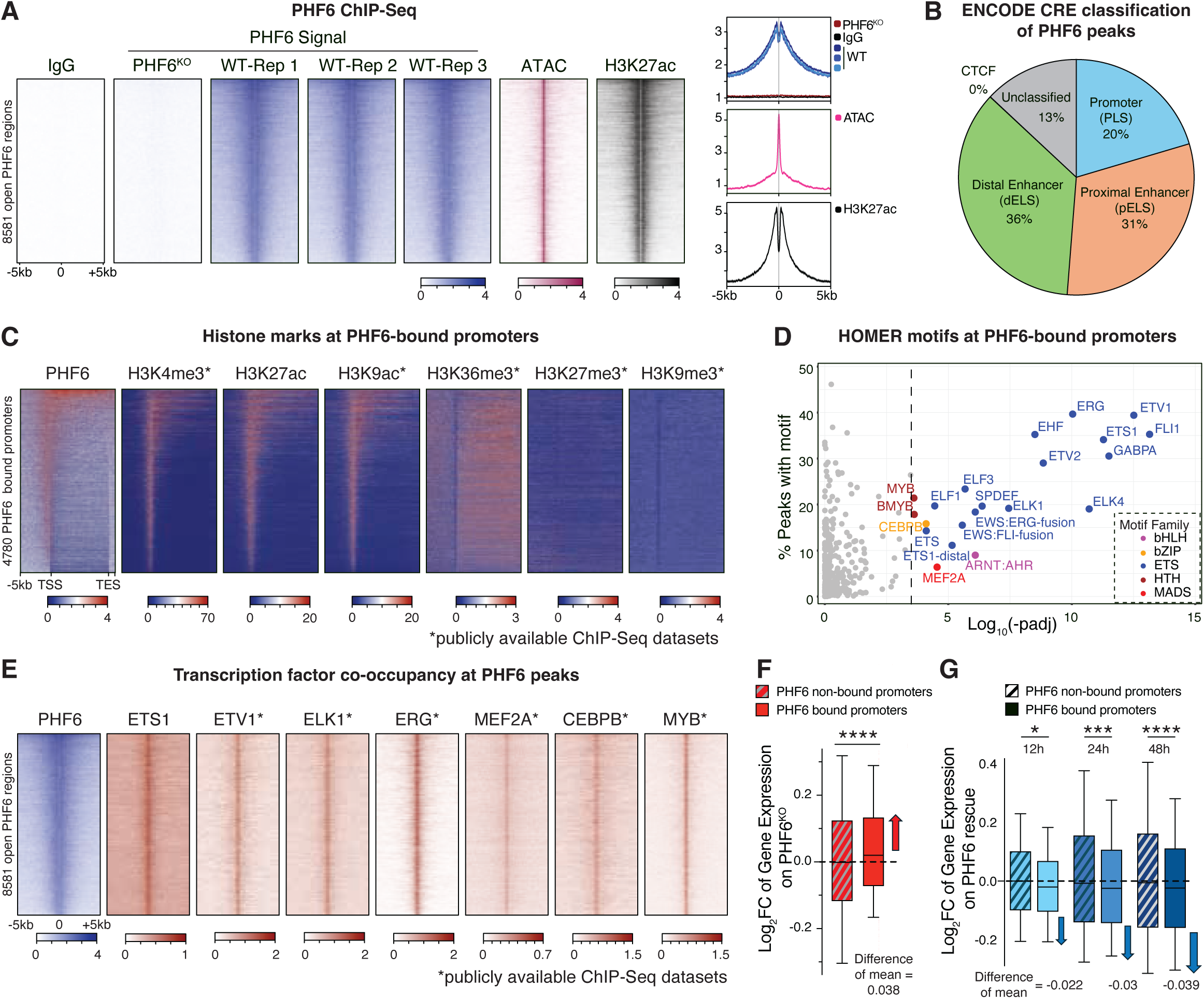
PHF6 binds gene promoters and represses transcription. A. Heatmaps (left) and meta-gene profiles (right) of 3 replicates of PHF6 ChIP-Seq signal at open high-confidence PHF6 peaks, along with ATAC-Seq and H3K27ac ChIP-Seq. IgG ChIP-Seq in WT and PHF6 ChIP-Seq in PHF6^KO^ are shown as negative controls. B. Pie chart showing categorization of PHF6 peaks based on overlap with ENCODE-defined cis-regulatory elements (CREs). C. Heatmaps showing PHF6 ChIP-Seq along with selected active and repressive histone modifications (from our lab and from publicly available datasets) along bodies of genes with PHF6-bound promoters. D. Scatter plot showing motifs and motif families enriched at PHF6-bound promoters. E. Heatmaps showing PHF6 co-occupancy with ETS family TFs, MEF2A, CEBPB, and MYB at PHF6-bound promoters. F. Boxplots showing differential expression in PHF6^KO^ compared to WT of genes with or without PHF6 binding at promoters. Boxplots show median (line), interquartile range (box), and minimum to maximum data range (whisker). G. Boxplots showing differential expression following time course of PHF6 rescue of genes with or without PHF6 binding at promoters. Boxplots show median (line), interquartile range (box), and minimum to maximum data range (whisker).

Motif analysis of PHF6-bound promoters showed a high degree of enrichment for ETS transcription factors (TFs), with a lower degree of enrichment for MEF, CEBP, and MYB motifs (**Fig** 2D). Alignment of publicly available ChIP-Seq tracks from hematopoietic or leukemic cell lines confirmed striking co-occupancy of PHF6 with ETS factors ETS1, ETV1, ELK1, and ERG (**Fig** 2E), in addition to co-occupancy with MEF2A, CEBPB and MYB (**Fig** 2E). To determine what effect PHF6 occupancy exerts on the transcription of bound promoters, we examined gene expression changes on *PHF6* knockout and rescue. Genes with PHF6 at promoters showed overall increased mRNA levels on *PHF6* knockout compared to non-bound promoters (**Fig** 2F). Reciprocally, the dox-induced rescue of PHF6 on a null background decreased their expression (**Fig** 2G). In contrast with a previous report^25^, PHF6 binding did not alter ATAC (**Fig** S2C) or H3K27ac (**Fig** S2D) signal at any PHF6-bound sites (promoters or non-promoters). Thus, in AML, PHF6 binds actively transcribed promoters bound by ETS factors and represses their transcription without any appreciable direct effect on chromatin accessibility or H3K27ac signal. This transcriptional repression effect is likely a small modulatory change distributed across a large set of bound genes.

### *R274Q* is a functionally null point mutation

To study the effects of *PHF6* missense mutations, we used CRISPR/Cas9-based homology-directed repair (HDR) to engineer *R274Q*, the most common *PHF6* missense mutation (**Fig** 1A), at the endogenous *PHF6* locus in THP-1 cells (**Fig** 3A, Top). Note: THP-1 has an XY genotype, with only one *PHF6* allele on its X chromosome. The resultant R274Q clones showed a 29% reduction in PHF6 protein levels (**Fig** 3A, bottom) without any change in mRNA levels (**Fig** 3B). Immunofluorescence imaging (IF) confirmed a small reduction in protein abundance without any change in its relative nucleoplasmic-nucleolar distribution (**Fig** 3C). Despite this modest reduction in protein, RNA-Seq showed that R274Q clones were transcriptionally similar to PHF6^KO^ clones (**Fig** 3D-E, S3A), with positive enrichment for HSC and progenitor genes (**Fig** 3F) and reduced expression of myeloid surface markers (**Fig** 3G).

**Figure 3:**
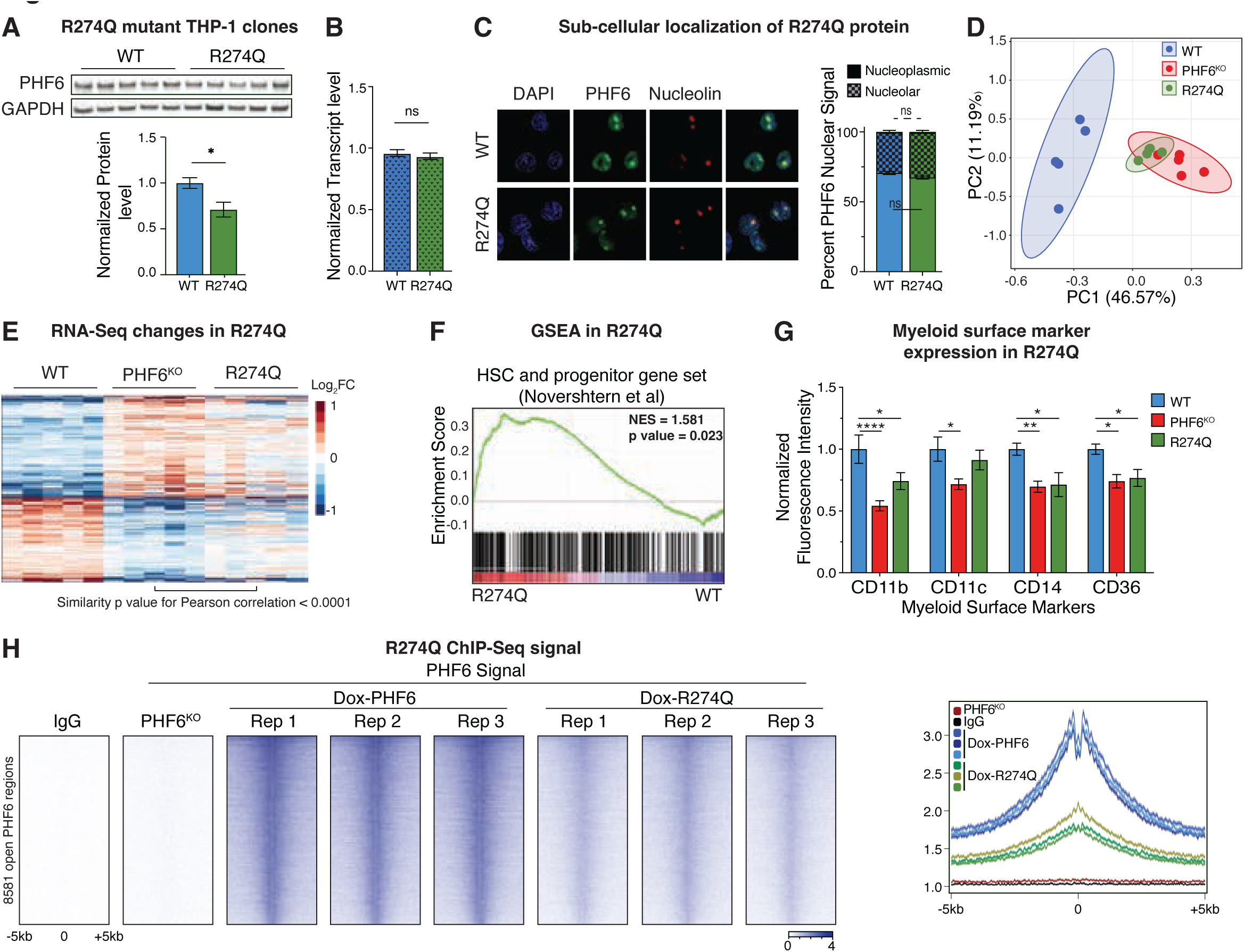
*R274Q* is a functionally null point mutation. A. Immunoblot (top) and bar graph showing quantification (bottom) of PHF6 protein in WT and R274Q clones in THP-1 cells. GAPDH is shown as loading control. (n=5) B. Bar graph showing RT-qPCR quantification of PHF6 mRNA levels in WT and R274Q. (n=3) C. Representative immunofluorescence images showing localization of PHF6 protein in WT and R274Q clones. DNA stain DAPI marks the nucleoplasm, and nucleolin is a nucleolar marker. Stacked bar graph shows distribution of PHF6 protein between nucleolus and nucleoplasm in WT and R274Q clones. (n=40-60 cells) D. Principal Component Analysis (PCA) plot of RNA-Seq replicates of WT, PHF6^KO^, and R274Q clones. E. Heatmaps showing effect of R274Q mutation on expression of genes differentially expressed in PHF6^KO^ compared to WT. Pearson correlation shows similarity between expression profiles of R274Q and PHF6^KO^ clones. F. GSEA plot showing positive enrichment of HSC and progenitor cell gene set in R274Q compared to WT. G. Bar graph showing normalized median fluorescence signal of myeloid surface markers in R274Q compared to WT, with PHF6^KO^ shown for comparison. (n=3) H. Heatmaps (left) and meta-gene profiles (right) of replicates of PHF6 and R274Q ChIP-Seq signal in doxycycline-inducible clones optimized to express identical levels of WT and mutant protein. PHF6 tracks are the same as those shown in Fig 2A. All bar graphs show mean ± standard error of mean (SEM), ns (not significant) = p ≥ 0.05, *p = 0.01 to 0.05, **p = 0.001 to 0.01, ***p = 0.001 to 0.0001, ****p < 0.0001, by one-way ANOVA with Sidak’s multiple comparison testing.

Since the transcriptional consequences of *R274Q* mutation seemed out of proportion to its protein level, we sought to examine whether R274Q protein has impaired ability to bind chromatin. Cognizant that reduced steady-state protein level may confound ChIP-Seq signal interpretation, we generated a dox-inducible clone expressing R274Q (Dox-R274Q) and used it in conjunction with a clone with dox-inducible wild-type PHF6 (Dox-PHF6). We optimized doxycycline doses to achieve similar levels of wild-type and mutant protein in a PHF6 knockout background (**Fig** S3B). ChIP-Seq with *Drosophila* spike-in chromatin normalization showed that R274Q protein, even when present within cells at levels matching wild-type PHF6, has consistently reduced chromatin occupancy at all sites (**Fig** 3H). RNA-Seq analysis of the inducible Dox-PHF6 and Dox-R274Q clones confirmed that while wild-type PHF6 induction on a null background was able to alter the expression of hundreds of genes, R274Q induction led to a virtual absence of any transcriptional change (**Fig** S3C). Collectively, our results show that *R274Q* is a functionally null mutation, with significantly impaired chromatin occupancy and a complete inability to exert downstream transcriptional effects.

### PHF6 missense mutations cause loss of function through compromised protein abundance and chromatin occupancy

Missense mutations of *PHF6* comprise one-third of its mutations in hematological malignancies^54^ (**Fig** 1A). Most of these missense mutations (including *R274Q*) are concentrated in the second extended PHD (ePHD2), suggesting that ePHD2 may play an important structural and/or functional role for the protein. To understand whether other ePHD2 mutations compromise PHF6 function like *R274Q* did, we conducted a detailed clinical and functional characterization of six ePHD2 missense mutations (*C242Y*, *D262V*, *R274Q*, *G287V*, *C297Y*, and *I314T*, chosen based on their frequency in hematopoietic malignancies) and three non-ePHD2 missense mutations (*C20G*, *P153S*, and *E340K*, falling within the ePHD1 domain, the region between ePHD1 and ePHD2, and the post-ePHD2 stretch respectively) (**Fig** 1A (right), 4A). In contrast to the ePHD2 mutations, which were recurrent, the non-ePHD2 mutations were each reported in only a single patient (**Fig** 4B).

**Figure 4:**
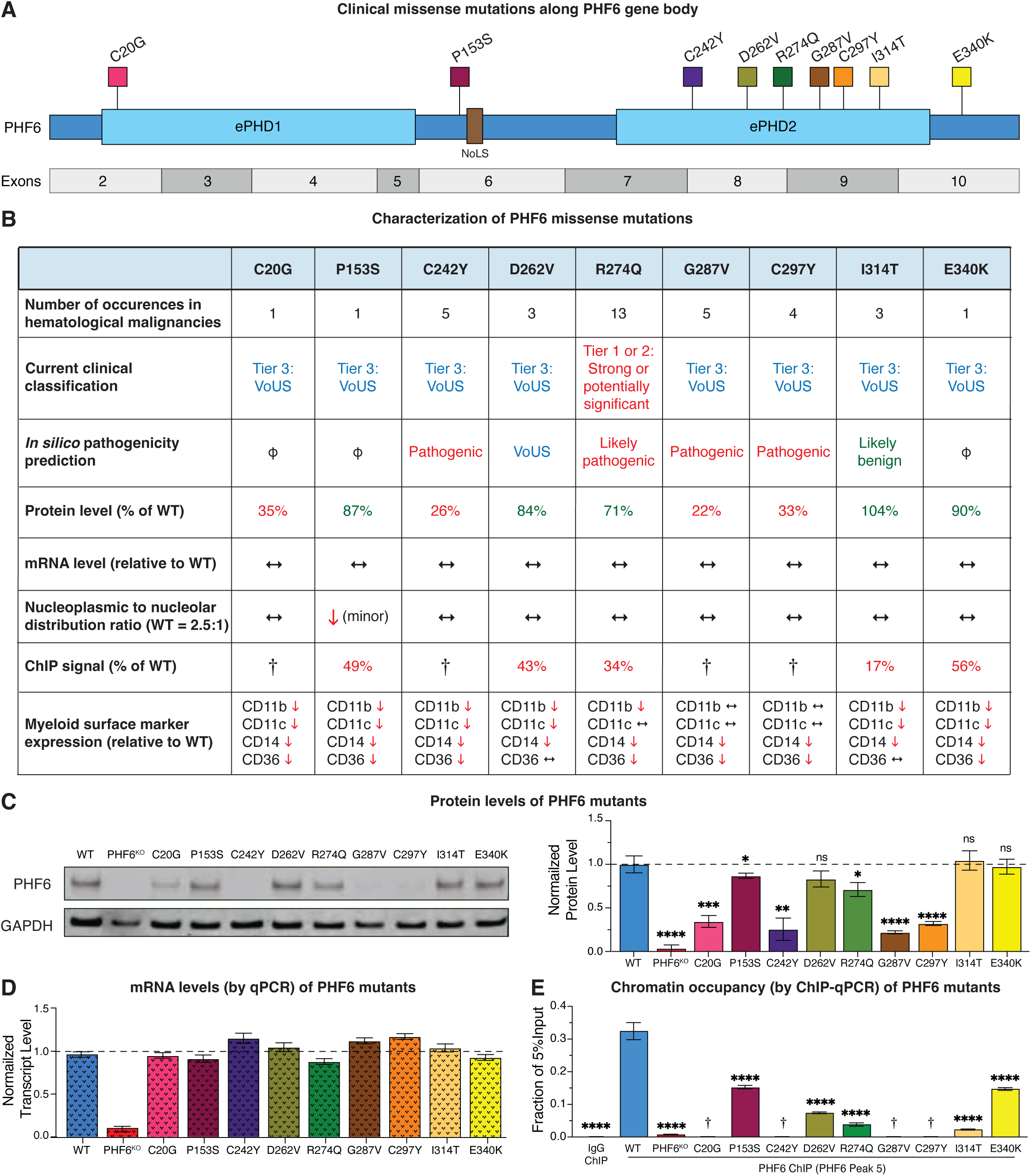
PHF6 missense mutations cause loss of function through compromised protein abundance and chromatin occupancy. A. Lollipop plot depicting nine PHF6 missense somatic mutations selected for functional dissection. C242Y, D262V, R274Q, G287V, C297Y and I314T are within the ePHD2 domain, while C20G, P153S and E340K are outside. NoLS: Nucleolar localization signal. B. Table summarizing functional characterization of PHF6 missense mutants (details in Fig S4). Patient numbers were obtained from COSMIC through the ProteinPaint portal. Clinical classification of mutations was performed by the Penn Center for Personalized Diagnostics. Pathogenicity prediction was performed on ePHD2 mutants using 4 concordant meta-predictors: REVEL, MetaLR, MetaSVM, and Condel (Fig S4D). ɸ indicates mutants unable to be analyzed due to the unavailability of structure for non-ePHD2 domains. For protein and mRNA levels, ↔ indicates no change. ChIP signal is the average ChIP-qPCR signal at 5 PHF6 peaks, † indicates mutants skipped for ChIP-qPCR due to low protein level. Values marked in red are considered pathogenic or functionally detrimental for the analysis in question. C. Immunoblots (left) showing PHF6 protein level in one representative clone for each missense mutation compared to WT and PHF6^KO^. GAPDH is shown as loading control. Bar graph (right) quantifies PHF6 protein in multiple replicate clones for each mutant (Fig 4SC), normalized to GAPDH. R274Q quantification shown here is the same as that shown in Fig 3A, and is included here for completeness. (n=4-9 clones for each mutant) D. Bar graph showing RT-qPCR quantification of PHF6 mRNA levels in mutant clones compared to WT. R274Q quantification shown here is the same as that shown in Figure 3B, and is included here for completeness. (n=3) E. Bar graph showing PHF6 ChIP-qPCR signal at a representative PHF6 peak in mutants compared to WT. (n=3), † indicates mutants skipped due to low protein levels (below 70% of WT clones). All bar graphs show mean ± standard error of mean (SEM), ns (not significant) = p ≥ 0.05, *p = 0.01 to 0.05, **p = 0.001 to 0.01, ***p = 0.001 to 0.0001, ****p < 0.0001, by one-way ANOVA with Sidak’s multiple comparison testing.

We began by having the Penn Center for Personalized Diagnostics (tasked with performing all clinical next-generation sequencing at the Hospital of the University of Pennsylvania) assign a classification to each mutation by running them through the standard clinical genomics workflow used for annotating variants identified in patient samples (**Fig** S4A). According to this workflow, only *R274Q* could be classified as a Tier 1 or 2 mutation (strong or potentially significant), while all others were classified as Tier 3 (VoUS - variant of unknown significance) (**Fig** 4B), highlighting the clinical challenges that arise in assessing pathogenicity of variants detected in patient biopsies. Next, we performed *in silico* analyses using a published crystal structure of the ePHD2 domain (amino acids 208-333)^22^, and applied four independent protein folding algorithms to predict the pathogenicity of each mutant based on its effects on structural stability (**Fig** S4B). These analyses predicted that *C242Y*, *R274Q*, *G287V*, and *C297Y* were pathogenic or likely pathogenic, while *D262V* was a VoUS, and *I314T* was likely benign (**Fig** 4B, S4B).

Next, we used CRISPR/Cas9-based HDR to engineer multiple clones of THP-1 cells with each mutation. PHF6 protein levels varied from low (22% to 35% of WT) in C20G, C242Y, G287V, and C297Y, to the 71% to 87% range in P153S, D262V, and R274Q, and to levels comparable to WT in I314T and E340K (**Fig** 4B-C & S4C). Immunofluorescence confirmed these reductions in protein levels, and showed no change in relative nucleoplasmic-nucleolar distribution, except in P153S, which had a ∼10% increase in the nucleolar fraction (**Fig** 4B, S4D-F). As expected, these variable protein levels were not caused by any change in *PHF6* mRNA levels (**Fig** 4B, 4D).

We next sought to determine whether any of the missense mutant proteins had reduced chromatin occupancy and myeloid surface marker expression similar to R274Q. ChIP-qPCR for all mutants with greater than 70% protein level compared to WT (P153S, D262V, R274Q, I314T, and E340K) showed variable levels of reduced chromatin occupancy out of proportion to their protein levels (**Fig** 4B, 4E, S4G). Finally, flow cytometry showed that all mutants showed reduced levels of two or more myeloid surface markers (**Fig** 4B, S4H). Collectively, our results show that though most *PHF6* missense mutations would currently be classified as VoUSs in the clinic, they uniformly cause compromised protein abundance, chromatin occupancy, or both.

### PHF6 cannot occupy chromatin without its functional partner PHIP, a newly-described AML-mutated protein

To identify possible functional partners for PHF6, we explored cancer cell line dependency data from the DepMap project^67,68^. We specifically examined the correlation of dependency scores, which range from 0 to 1 for every possible pair of genes, depending on how similar the effects of CRISPR knockouts of the two genes are on the growth of 1,150 cell lines screened as part of DepMap. The gene with the highest correlation of dependency with *PHF6* was *PHIP* (Pleckstrin Homology domain Interacting Protein), with a correlation score of 0.54 (**Fig** 5A-B). PHIP (also known as DCAF14, RepID, BRWD2) is one of many substrate receptor proteins for the cullin4A-RING E3 ubiquitin ligase (CRL4) complex^77^. Examination of the literature revealed multiple additional lines of evidence supporting a common biological role for PHF6 and PHIP. PHF6 was among the proteins identified by mass spectrometry in a chromatin immunoprecipitation study of PHIP in HEK-293 cells^78^. Both genes are recurrently mutated in acute and chronic myeloid malignancies^1,11,32,79–82^ (**Fig** 5C), and though *PHIP* is mutated in <1% in most databases, a recent study of 100 Black patients with AML identified *PHIP* mutations in 7 cases^32^. Most provocatively, germline mutations in *PHF6* and *PHIP* produce rare neurodevelopmental syndromes^83–86^ with highly overlapping clinical features^87–89^ (**Fig** 5D), providing a strong basis to investigate whether the two proteins may be performing a unified biological function.

**Figure 5:**
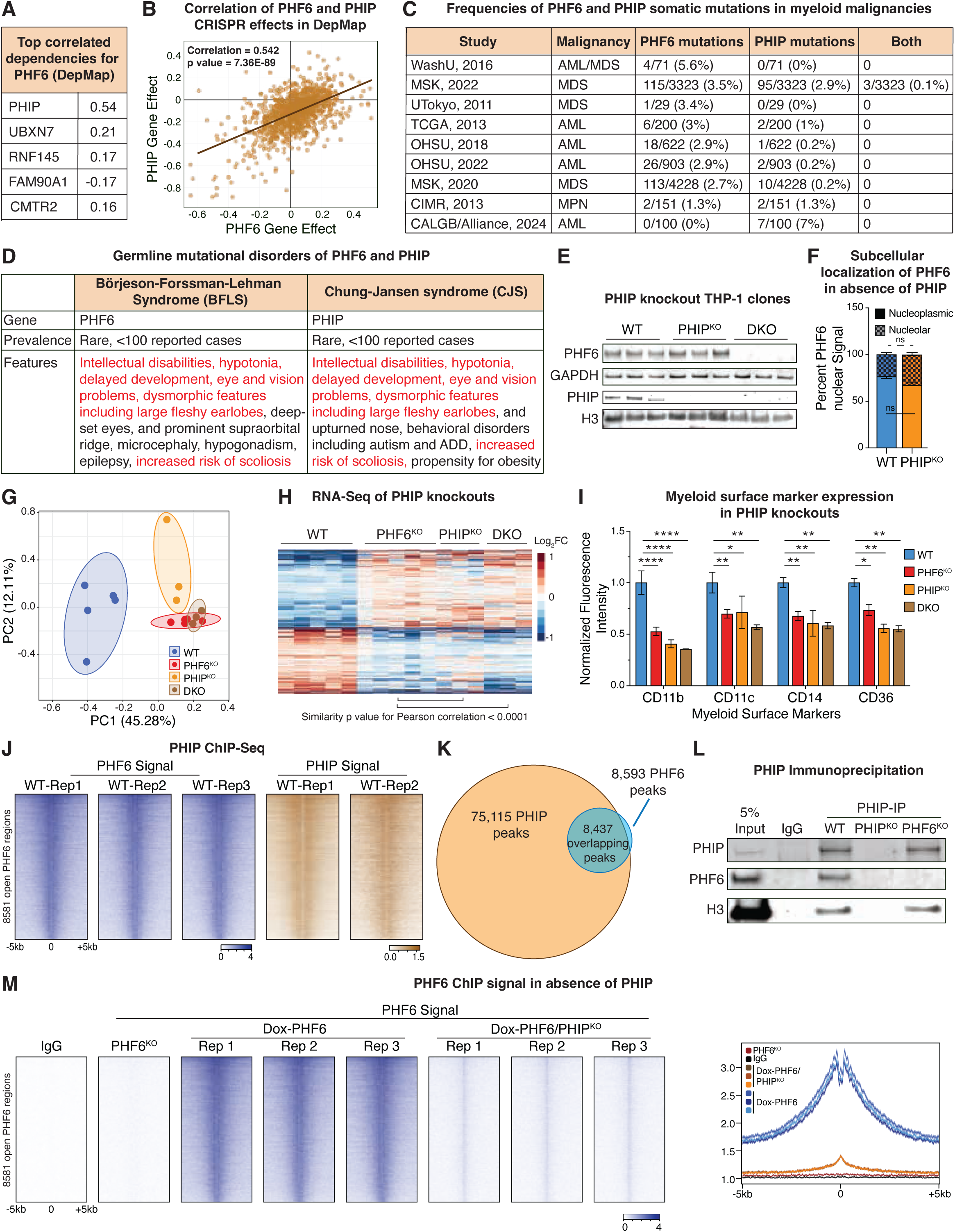
PHF6 cannot occupy chromatin without its functional partner PHIP, a newly-described AML-mutated protein. A. Table showing the top five correlated gene dependencies for *PHF6* in the Broad Institute DepMap project. B. Scatter plot showing correlation of CRISPR screen (Chronos) gene scores for *PHF6* and *PHIP* in 1,150 cell lines screened in DepMap. C. Table showing frequencies of *PHF6* and *PHIP* mutations in databases of patients with myeloid neoplasms. Data were obtained from cBioPortal. D. Table summarizing features of rare neurodevelopmental syndromes caused by germline mutations of *PHF6* and *PHIP*. Features marked in red are common between both syndromes. E. Immunoblots of PHF6 and PHIP in PHIP^KO^ and DKO clones. GAPDH and H3 are shown as loading controls. F. Stacked bar graph showing distribution of PHF6 protein between nucleolus and nucleoplasm in WT and PHIP^KO^ clones. (n=40-60 cells) G. PCA plot of RNA-Seq replicates of WT, PHF6^KO^, PHIP^KO^ and DKO clones. H. Heatmap showing effects in PHIP^KO^ and DKO on expression of genes differentially expressed in PHF6^KO^ compared to WT. Pearson correlation shows similarities between expression profiles of single and double knockout clones. I. Bar graph showing normalized median fluorescence signal of myeloid surface markers in PHIP^KO^ and DKO clones compared to WT clones. (n=3) J. Heatmaps showing PHIP ChIP-Seq signal at PHF6 peaks. PHF6 tracks are the same as those shown in Fig 2A and Fig 3H. K. Venn diagram showing overlap of PHF6 and PHIP peaks. L. Immunoblots showing pulldown of PHF6 with PHIP-ChIP. IgG-ChIP, and PHIP-ChIP in PHIP^KO^ and PHF6^KO^ clones are shown as negative controls. H3 is shown as a positive control for chromatin pulldown. M. Heatmaps (left) and meta-gene profiles (right) of replicates of PHF6 ChIP-Seq signal in WT and PHIP^KO^ clones. PHF6 tracks are the same as those shown in Fig 2A, 3H, and 3J. All bar graphs show mean ± standard error of mean (SEM), ns (not significant) = p ≥ 0.05, *p = 0.01 to 0.05, **p = 0.001 to 0.01, ***p = 0.001 to 0.0001, ****p < 0.0001, by one-way ANOVA with Sidak’s multiple comparison testing.

To study whether PHF6 and PHIP are mechanistically linked in AML, we generated single-knockout clones of *PHIP* (PHIP^KO^) as well as double-knockout clones of *PHF6* and *PHIP* (DKO) in the THP-1 line (**Fig** 5E). Using immunoblotting and immunofluorescence, we observed that *PHIP* knockout does not affect PHF6 protein level (**Fig** 5E, S5A) or nucleoplasmic-nucleolar distribution (**Fig** 5F, S5B). RNA-Seq of PHIP^KO^ and DKO clones showed their striking transcriptional similarity to PHF6^KO^ clones (**Fig** 5G-H, S5C-D), with each showing positive enrichment for HSC and progenitor genes (**Fig** S5E), and reduced myeloid cell surface markers identical to PHF6^KO^ (**Fig** 5I).

To determine whether PHF6 and PHIP have overlapping chromatin occupancy, we performed ChIP-Seq for PHIP and observed that while PHIP showed more ChIP-Seq peaks (75,115) than PHF6 (8,593), ∼98% of PHF6 peaks overlapped with PHIP peaks, with strong alignment of occupancy patterns (**Fig** 5J-K). Two-thirds of PHIP peaks showed ATAC and H3K27ac ChIP signal (**Fig** S5F-G). Interestingly, PHIP peaks with open chromatin (those overlapping with ATAC peaks) showed higher ChIP-Seq signal for both PHIP and PHF6 (**Fig** S5H) compared to PHIP peaks with closed chromatin. ChIP-immunoblotting of PHIP confirmed pulldown of PHF6 (**Fig** 5L). To examine whether PHIP loss affects the ability of PHF6 to occupy chromatin, we generated *PHIP* knockout clones in the doxycycline-inducible system (Dox-PHF6/PHIP^KO^) (**Fig** S5I, left). PHIP loss had no effect on the rescue of PHF6 protein levels, (**Fig** S5I, Right), but Dox-PHF6/PHIP^KO^ clones showed a near complete loss of PHF6 chromatin occupancy (**Fig** 5M). RNA-Seq analysis of the doxycycline-inducible system showed that the downstream transcriptome changes produced on PHF6 induction were blunted in the absence of PHIP (**Fig** S5J). Collectively, our results show that the chromatin occupancy and downstream transcriptional effects of PHF6 are dependent on its functional partner PHIP.

## Discussion

Our studies show that PHF6 is a transcriptional repressor that binds to open and active promoters and enhancers, and either directly or indirectly represses a stemness gene network in AML (**Fig** 6A). Our comprehensive dissection of multiple clinical PHF6 missense mutations shows that all assayed mutants are either hypomorphic or functionally null due to a combination of reduced protein abundance and compromised chromatin occupancy. Our work also reveals a novel mechanistic connection between PHF6 and PHIP, with the former depending on the latter for its chromatin occupancy, and their knockouts producing similar downstream transcriptional and surface marker effects. Through this work, we provide evidence that these two disparate leukemia-mutated proteins suppress stemness as part of the same functional chromatin complex.

**Figure 6:**
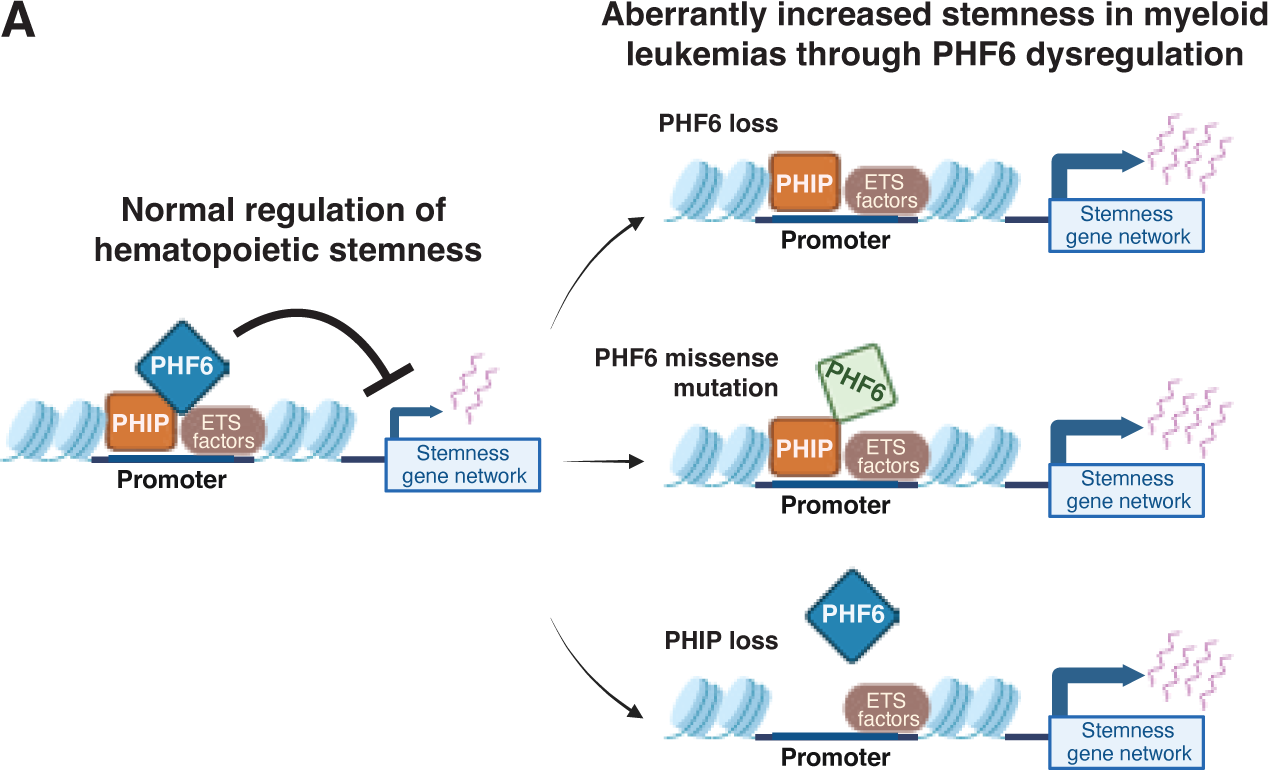
PHF6-PHIP complex represses AML stemness. A. Model of PHIP-dependent PHF6 role in hematopoietic and leukemic stemness: PHF6 and PHIP form a complex on promoters bound by ETS factors and repress their transcription, thereby repressing a stemness gene network. In a subset of acute or chronic myeloid malignancies, loss of PHF6 chromatin occupancy, either through loss of PHF6 itself, through missense mutations in PHF6 that impair its protein stability or chromatin occupancy, or through loss of PHIP, eliminates this repression and increases stemness.

*PHF6* mutations are associated with worse prognosis in MDS and AML^26,90^. Accurately classifying mutations in the clinic is critical for prognostication and medical decision-making. Missense variants, especially in understudied genes, often end up classified as VoUSs in patient reports, posing challenges for oncologists attempting to interpret next-generation sequencing results. An example of this challenge is the fact that a standard clinical workflow classified 8 of 9 mutations we assayed as VoUSs, and *in silico* analyses could only recognize 4 as pathogenic. Engineering and characterization of mutations in the THP-1 AML cell line, which we have shown in this work to be a reliable model to study PHF6, revealed that all 9 of them impaired PHF6 function, with some leading to reduced protein abundance (C20G, C242Y, G287V, C297Y), and others largely sparing protein levels but acting through compromised chromatin occupancy (P153S, D262V, R274Q, I314T, E340K). None of the mutations caused any meaningful change in the nucleoplasmic-nucleolar distribution of PHF6 protein, though P153S did produce a ∼10% shift from the nucleoplasmic fraction to the nucleolus. We speculate that this shift could have been caused by the proximity of the mutation to a predicted nucleolar localization signal^22^, though it is implausible that such a minor change could account for the magnitude of functional impairment of the mutant. Future work to identify mutations that completely uncouple nucleolar and nucleoplasmic localization of PHF6 would nonetheless be of value to dissect which fraction of the protein is primarily responsible for its stemness suppressive function.

We note that *in silico* analyses, which were informed by a previously published PHF6 ePHD2 domain crystal structure, assigned deleterious scores to ePHD2 mutants that turned out to have significant reductions in protein levels. This demonstrates the ability of such tools to recognize amino acid substitutions that produce structural clashes significant enough to impair protein stability. However, substitutions that disproportionately affected chromatin occupancy could not be predicted, indicating that these residues are likely responsible not for the internal stability of PHF6 protein, but for external contacts with chromatin interactors, potentially including PHIP. Future work will be required to determine whether PHF6 directly interacts with PHIP, and whether residues like R274 are part of the interaction surface.

*PHIP* recently came to attention in myeloid malignancies when mutations in it were identified in a small but meaningful subset of Black patients with AML (7/100 cases), compared to <1% cases in databases that had focused largely on White patients^32^. Current mutational numbers are too sparse to statistically quantify the correlation or anti-correlation of *PHF6* and *PHIP* mutations. Our work showing that the two proteins suppress stemness as a common functional complex suggests that their mutations will likely be redundant and therefore anti-correlated. Our work also provides clues for future investigation into the collective function of the PHF6-PHIP complex in hematological malignancies. PHIP (also known in the literature as DCAF14, RepID, and BRWD2) belongs to a class of DCAF (DDB1-CUL4-Associated Factors) proteins that determine the substrate specificity of the cullin4A-RING E3 ubiquitin ligase (CRL4) complex. There is little consensus on the role of this particular DCAF, and the few groups that have studied it in non-hematopoietic cells have reported a diversity of possible functions. PHIP has been reported to guide CRL4 to place a non-degradative ubiquitination mark on the histone chaperone FACT complex, and this ubiquitination mark regulates the loading of histones onto freshly replicated DNA^91^. Alternatively, more complex mechanisms have been proposed involving competition of PHIP with other DCAFs to protect replication forks and mitotic spindle assembly^92–94^. Such mechanisms would likely not explain the strong co-occupancy we observe of PHF6 and PHIP at promoters and enhancers (in patterns matching active histone marks), nor would they plausibly explain the dynamic and reversible gene expression changes produced by *PHF6* knockout and rescue (also recapitulated by *PHIP* knockout). On the other hand, a publication that previously noted PHF6 among the proteins pulled down on PHIP immunoprecipitation reported that PHIP occupancy strongly correlated with H3K4me3, H3K4me1, and H3K27ac marks^78^. The same paper reported that a cryptic Tudor domain in PHIP was required for its ability to occupy chromatin and interact with methylated H3K4 histone peptides. Based on this latter report, we speculate that in hematopoietic and leukemic stem cells, PHIP binds active chromatin through direct recognition of histone modifications by its Tudor domain, and recruits PHF6. The complex of the two then potentially acts as a combined adapter complex to bridge the E3 ligase CRL4 to ubiquitinate a currently unknown chromatin target. Testing whether PHIP recruits the rest of the CRL4 complex to chromatin, and identifying chromatin proteins in AML showing reduced ubiquitination in the absence of PHF6 or PHIP, would be critical next steps to testing this model and determining how these two leukemia-mutated proteins function as a unified chromatin complex to repress hematopoietic and leukemic stemness.

## Acknowledgments

We thank Nancy Speck, Ivan Maillard, Wei Tong, Kathrin Bernt, Liling Wan, and Alessandro Gardini for helpful discussions. VRP is supported by National Institute of Health (NIH) grant R01-HL155144 (NHLBI), R35-GM138035 (NIGMS), American Cancer Society (ACS) grant 129784-IRG-16-188-38-IRG, an American Society of Hematology (ASH) Faculty Scholar Award, and the University of Pennsylvania Covid-19 Research Disruption Mitigation Fund. ASP is a Cell and Molecular Biology Student through the Biomedical Graduate Studies program at the University of Pennsylvania. CA is supported by a Fellow Scholar Award from the American Society of Hematology (ASH) and Co-Operative Center for Excellence in Hematology (CCEH) Type B grant by the National Institute of Diabetes and Digestive and Kidney Diseases (NIDDK). KFL is funded by NIH grants R35-GM133721 (NIGMS) and R01-HL171617. Research on this project was supported by the UPenn Abramson Cancer Center Support Grant, supported by NCI P30 CA016520.

## Authorship Contributions

VRP conceived the project and supervised the study. ASP designed and performed the majority of experiments, with additional studies by PS, AA, CA, and RV. SSG and ASP performed bioinformatic analyses, including writing scripts for graphical representation. ASP made and edited figures. SKWB and JM performed clinical classification of PHF6 variants. ASP, AA, and MK performed imaging analysis, and MSMF and KFL assisted with microscopy and imaging. ASP and VRP wrote the manuscript with input from all authors.

