## Supplemental Figures for "Leukemia-mutated proteins PHF6 and PHIP form a chromatin complex that represses acute myeloid leukemia stemness"

### Supplementary Figure 1

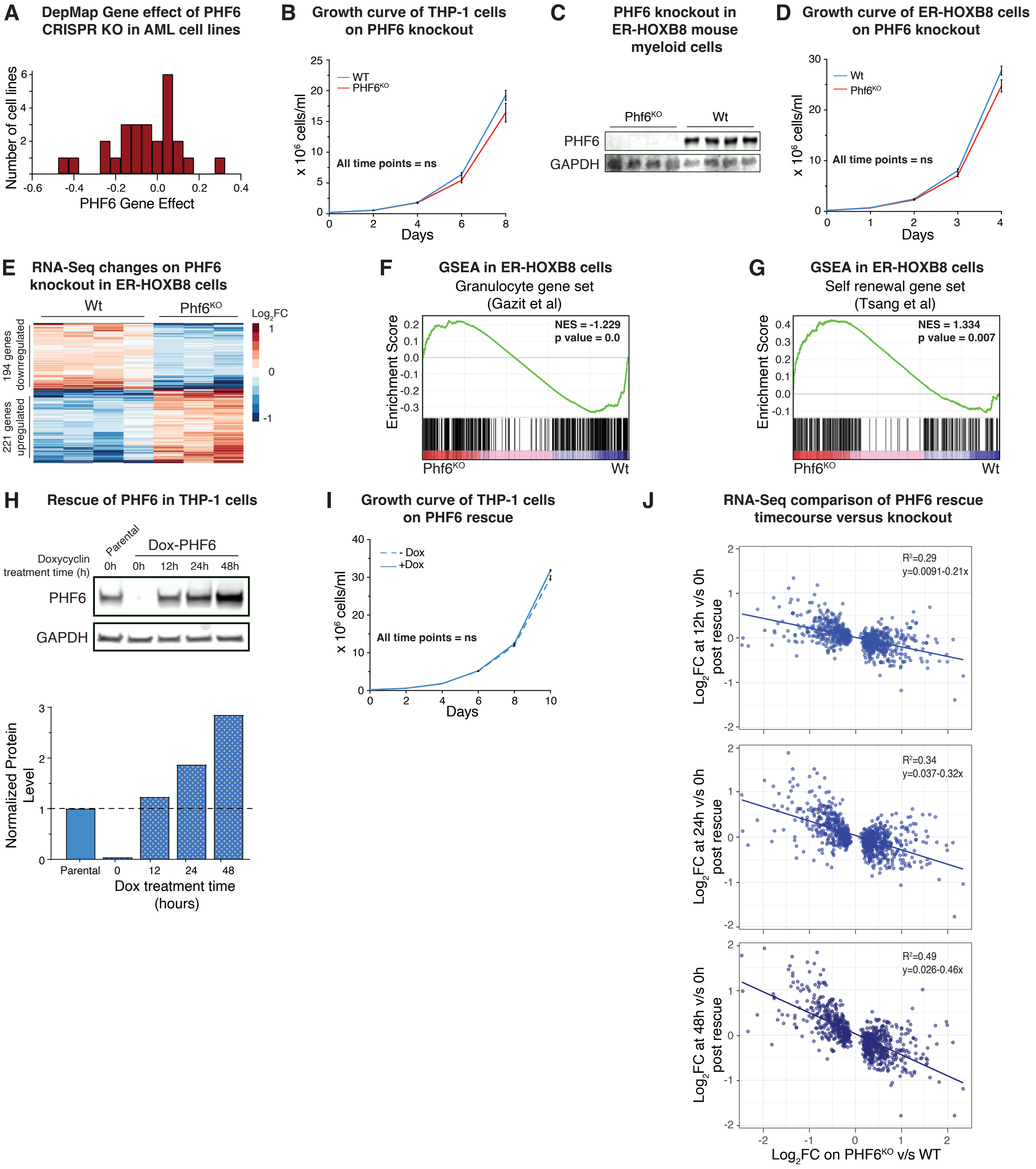

**Figure S1 (related to Fig 1)**

A. Histogram showing Chronos gene effect scores (representing proliferation relative to negative controls) for *PHF6* CRISPR knockout in 26 AML cell lines profiled by the DepMap project.

B. Growth curves for WT and $PHF6^{KO}$ human THP-1 clones. (n=7)

C. Immunoblot for PHF6 in Wt and $Phf6^{KO}$ clones of the mouse ER-HOXB8 cell line. GAPDH is shown as loading control.

D. Growth curve for Wt and $Phf6^{KO}$ mouse ER-HOXB8 clones. (n=4)

E. Heatmap showing 412 differentially expressed genes in $Phf6^{KO}$ ER-HOXB8 clones compared to Wt.

F. GSEA plot showing negative enrichment of granulocyte gene set in $Phf6^{KO}$ ER-HOXB8 clones compared to Wt.

G. GSEA plot showing positive enrichment of self-renewal gene set in $Phf6^{KO}$ ER-HOXB8 clones compared to Wt.

H. Top, Immunoblot showing time course of rescue (overexpression) of PHF6 using 250 ng/ml doxycycline in the Dox-PHF6 subclone (THP-1 $PHF6^{KO}$ null background). GAPDH is shown as loading control. Bottom, Quantification of PHF6 protein rescue in Dox-PHF6 cells, normalized to GAPDH.

I. Growth curve of Dox-PHF6 cells in presence and absence of doxycycline (250 ng/ml). (n=3)

J. Scatter plots depicting comparative effects of *PHF6* knockout and PHF6 rescue on genes found to be differentially expressed in $PHF6^{KO}$ compared to WT. Negative correlation is observed between changes produced on *PHF6* knockout and those produced by time course PHF6 rescue (12, 24, and 48 hrs compared to 0 hr).

All bar graphs show mean ± standard error of mean (SEM), ns (not significant) = $p \geq 0.05$, *p = 0.01 to 0.05, **p = 0.001 to 0.01, ***p = 0.001 to 0.0001, ****p < 0.0001, by one-way ANOVA with Sidak's multiple comparison testing.

#### Supplementary Figure 2

**A** **Epigenetic marks and PHF6 peaks at ENCODE-defined CREs**

| | H3K4me3 Signal | H3K27ac Signal | TSS proximity | ATAC Signal | CTCF Signal | PHF6 Peaks |
|---|---|---|---|---|---|---|
| **Promoter-like Signature (PLS)** | Yes | | < 200bp | Yes | | 1754/8593 |
| **Proximal enhancer-like Signature (pELS)** | | Yes | < 2000bp | Yes | | 2649/8593 |
| **Distal enhancer-like Signature (dELS)** | | Yes | > 2000bp | Yes | | 3068/8593 |
| **CTCF element** | | | | Yes | Yes | 2/8593 |
| **Unclassified open chromatin** | Yes | | | Yes | | 1108/8593 + 12 non-overlapping |

**B** **PHF6 ChIP-Seq signal on rDNA**

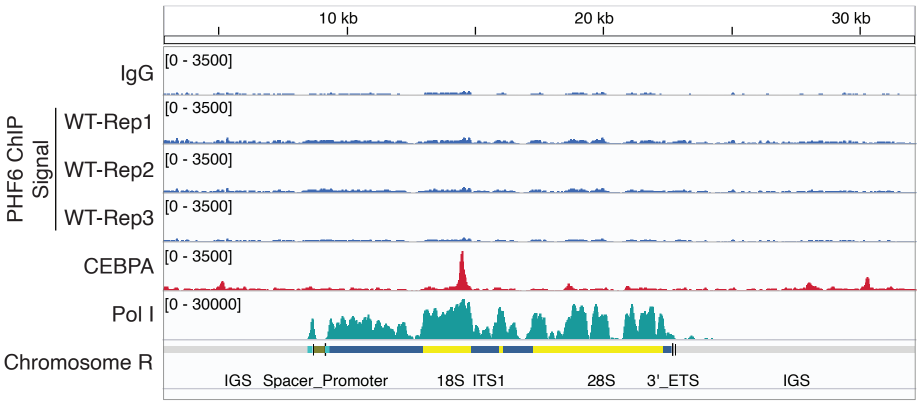


**C** ATAC Signal

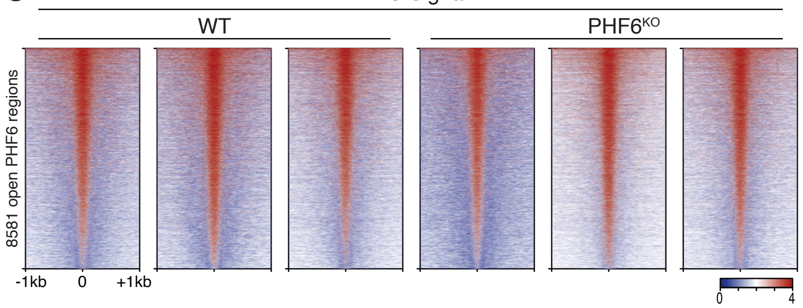


**D** H3K27ac ChIP Signal

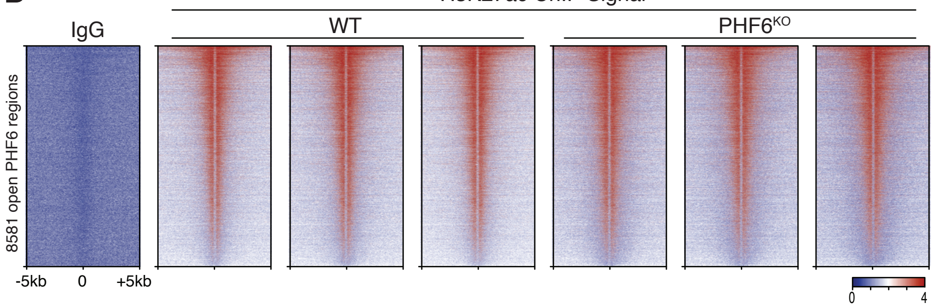

**Figure S2 (related to Fig 2)**

A. Table showing epigenetic marks used by ENCODE to define CREs, and the distribution of PHF6 peaks among these categories.

B. IGV genome browser tracks showing lack of alignment of PHF6 ChIP-signal (n=3) at ribosomal DNA. CEBPA and RNA Polymerase I (Pol I) alignment are shown as positive controls for rDNA mapping.

C. Heatmap of ATAC-Seq signal in WT and PHF6$^{KO}$ clones at PHF6-occupied open chromatin regions.

D. Heatmap of H3K27ac ChIP-Seq signal in WT and PHF6$^{KO}$ clones at PHF6-occupied open chromatin regions.

#### Supplementary Figure 3

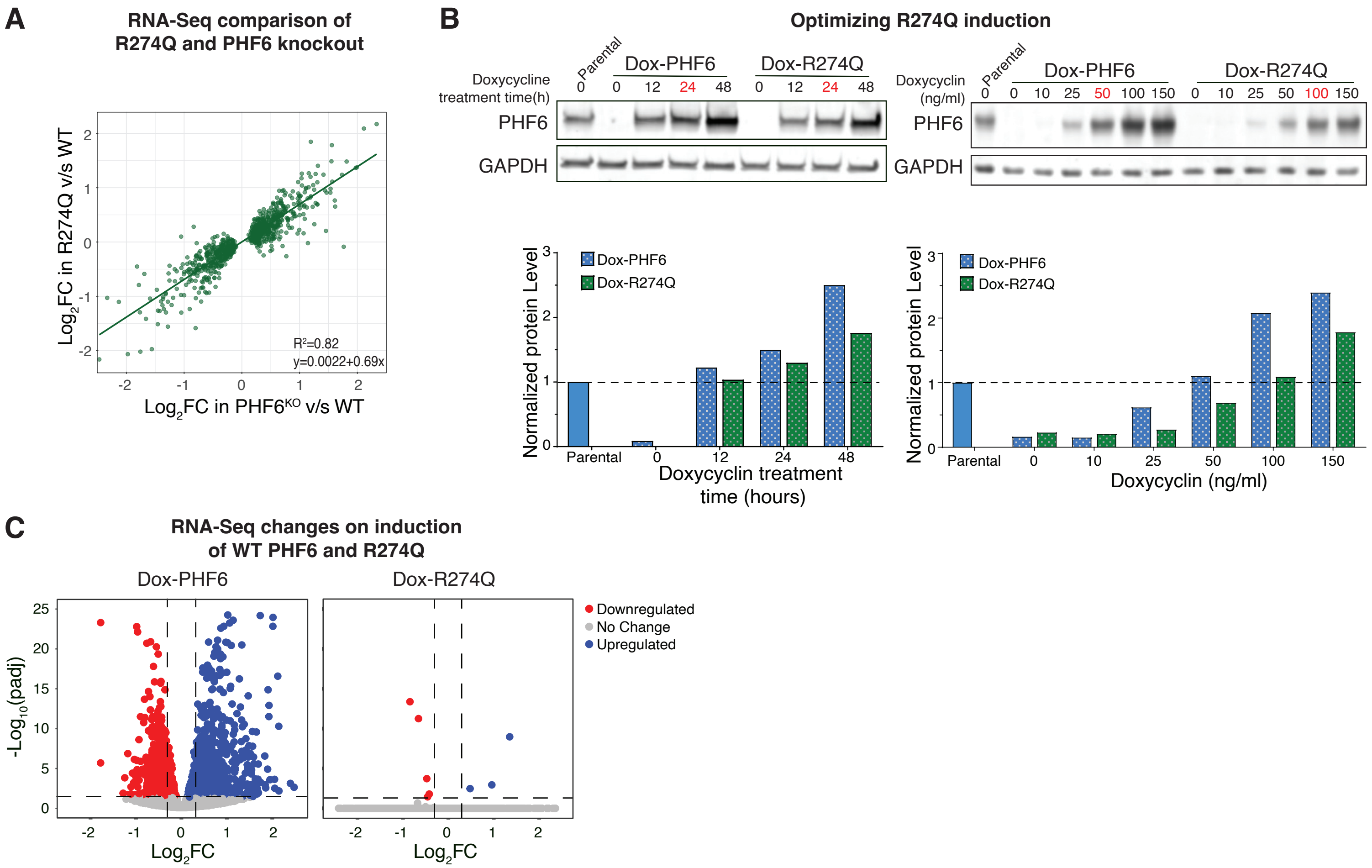

**Figure S3 (related to Fig 3)**

A. Scatter plot showing positive correlation between effects of *PHF6* knockout and *R274Q* mutation on genes found to be differentially expressed in $PHF6^{KO}$ compared to WT.

B. Immunoblots (top) and bar graphs (bottom) showing time and dose-dependent rescue of PHF6 in Dox-PHF6 and Dox-R274Q clones. GAPDH is shown as loading control and used for normalization. Time and dose used to induce PHF6 and R274Q for ChIP-Seq (in order to achieve equal protein levels of wild-type PHF6 and R274Q protein) are marked in red.

C. Volcano plots showing differential expression of genes on PHF6 rescue (overexpression) in Dox-PHF6 and Dox-R274Q clones.

### Supplementary Figure 4

**A** Tiers of clinical characterization used by the Penn Center for Personalized Diagnostics

| Tier | Categorization |
|---|---|
| Tier 1 | FDA approved therapy associated with the variant |
| Tier 1 or 2 | Pathogenic (disease-associated) or likely pathogenic variant without directed therapy |
| Tier 3 | VoUS, likely benign or benign varaints not reported |

**B** *In silico* analysis PHF6 missense mutants

| | C242Y | D262V | R274Q | G287V | C297Y | I314T |
|---|---|---|---|---|---|---|
| REVEL Score | 0.957 | 0.598 | 0.591 | 0.905 | 0.968 | 0.325 |
| Meta SVM Score | 1.002 | 0.4324 | 0.9127 | 0.844 | 1.0896 | -0.9988 |
| Meta LR Score | 0.8858 | 0.6519 | 0.8655 | 0.8214 | 0.991 | 0.1477 |
| Condel Score | 0.682 | 0.507 | 0.640 | 0.704 | 0.678 | 0.499 |

**C** PHF6 missense mutant clones

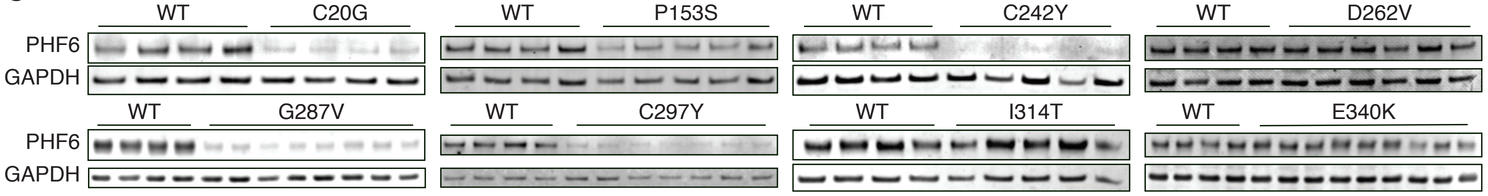


**D** Localization of PHF6 missense mutants

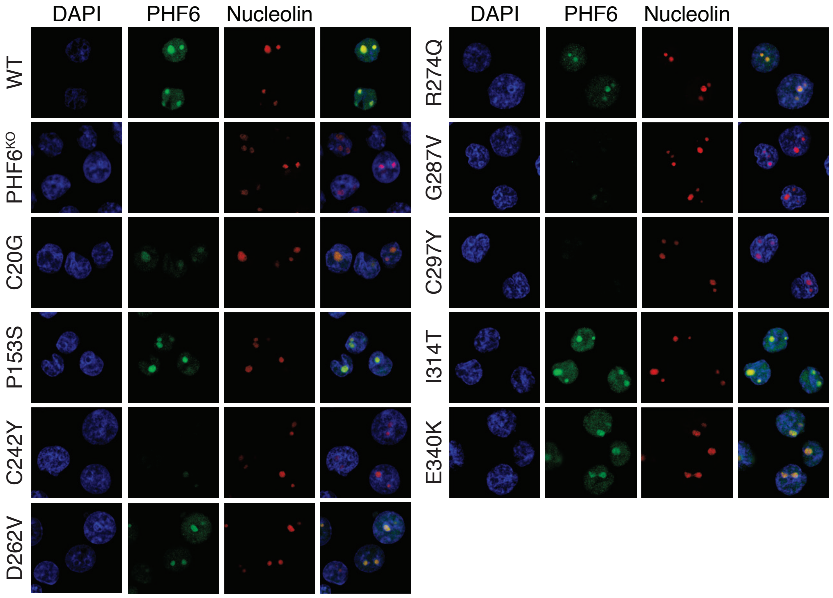


**E** Total nuclear signal of PHF6 missense mutants

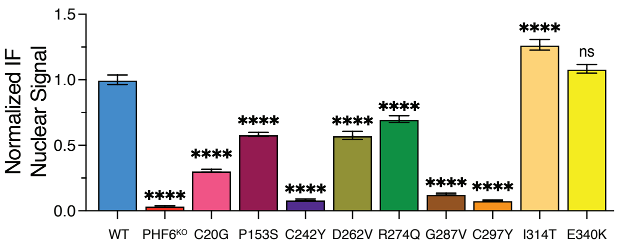


**F** Localization of PHF6 missense mutants

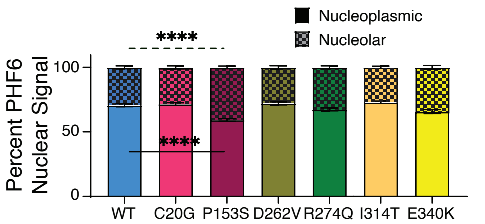


**G** PHF6 ChIP signal in PHF6 missense mutants

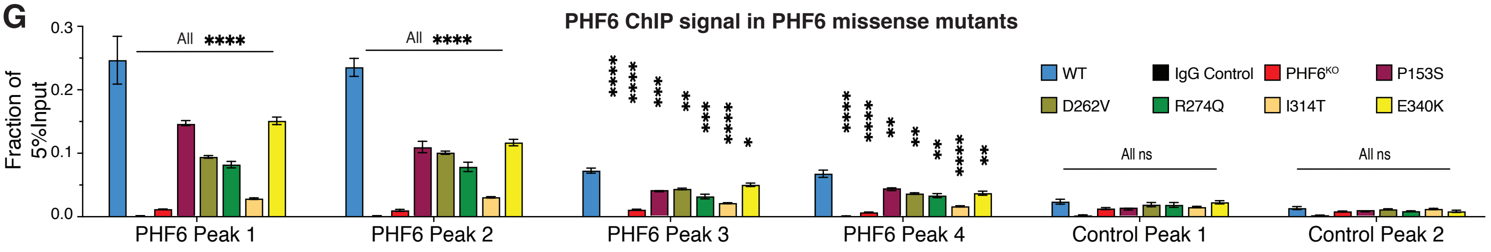


**H** Myeloid surface marker expression in PHF6 missense mutants

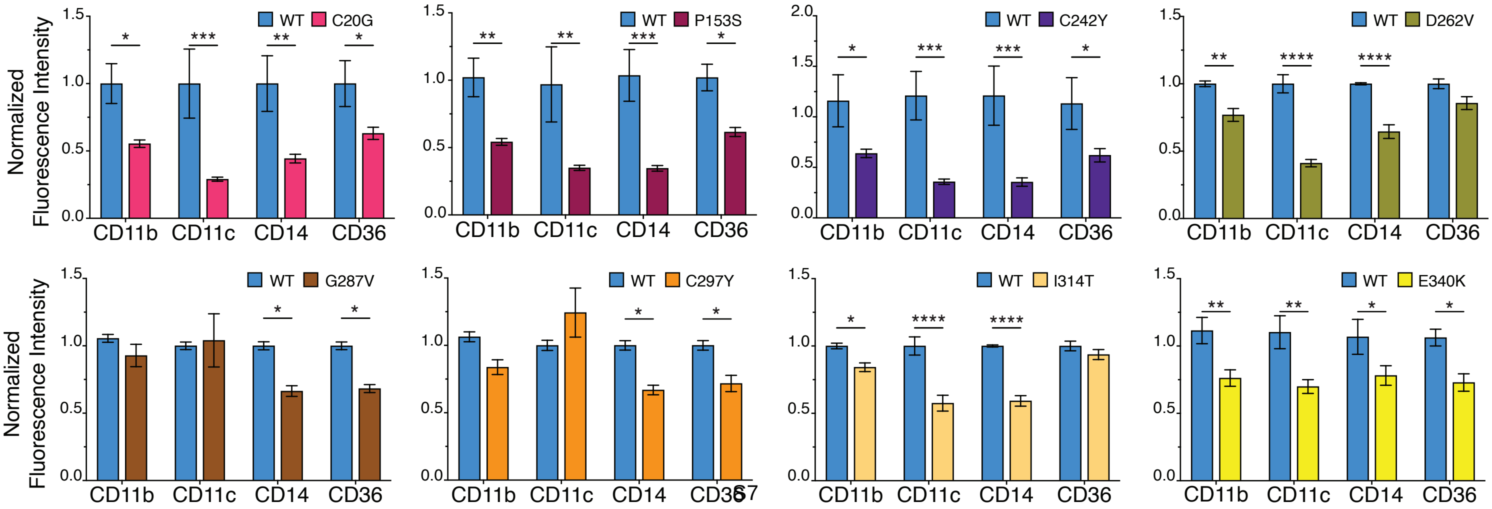

**Figure S4 (related to Fig 4)**

A. Table listing tiers of clinical classification used by the Penn Center of Personalized Diagnostics in its clinical genomics workflow applied to patient samples.

B. Table showing REVEL, Meta SVM, Meta LR, and Condel scores for ePHD2 mutations of PHF6. Scores marked in red are pathogenic and those marked in green are considered benign according to the scoring system for each prediction algorithm.

C. Immunoblots of PHF6 in PHF6 missense mutant clones. GAPDH is shown as loading control.

D. Representative immunofluorescence images showing localization of PHF6 protein in WT, $PHF6^{KO}$, and PHF6 missense mutants. DNA stain DAPI marks the nucleoplasm and nucleolin is a nucleolar marker.

E. Bar graph showing quantification of normalized total nuclear immunofluorescence signal for PHF6 in WT, $PHF6^{KO}$, and PHF6 missense mutant clones. (n=30-60 cells)

F. Stacked bar graph showing the distribution of PHF6 protein between nucleolus and nucleoplasm in WT and missense mutant clones with medium or high steady-state expression levels of PHF6. R274Q quantification is the same as Fig 3C and is included here for completeness. (n=40-60 cells)

G. Bar graphs showing PHF6 ChIP-qPCR signal at four PHF6 peaks and two control peaks in PHF6 missense expression mutants compared to WT clones as a fraction of 5% input. (n=3)

H. Bar graphs showing normalized expression of myeloid surface markers in PHF6 missense mutant clones compared to WT clones. (n=3)

All bar graphs show mean ± standard error of mean (SEM), ns (not significant) = p ≥ 0.05, *p = 0.01 to 0.05, **p = 0.001 to 0.01, ***p = 0.001 to 0.0001, ****p < 0.0001, by one-way ANOVA with Sidak's multiple comparison testing.

#### Supplementary Figure 5

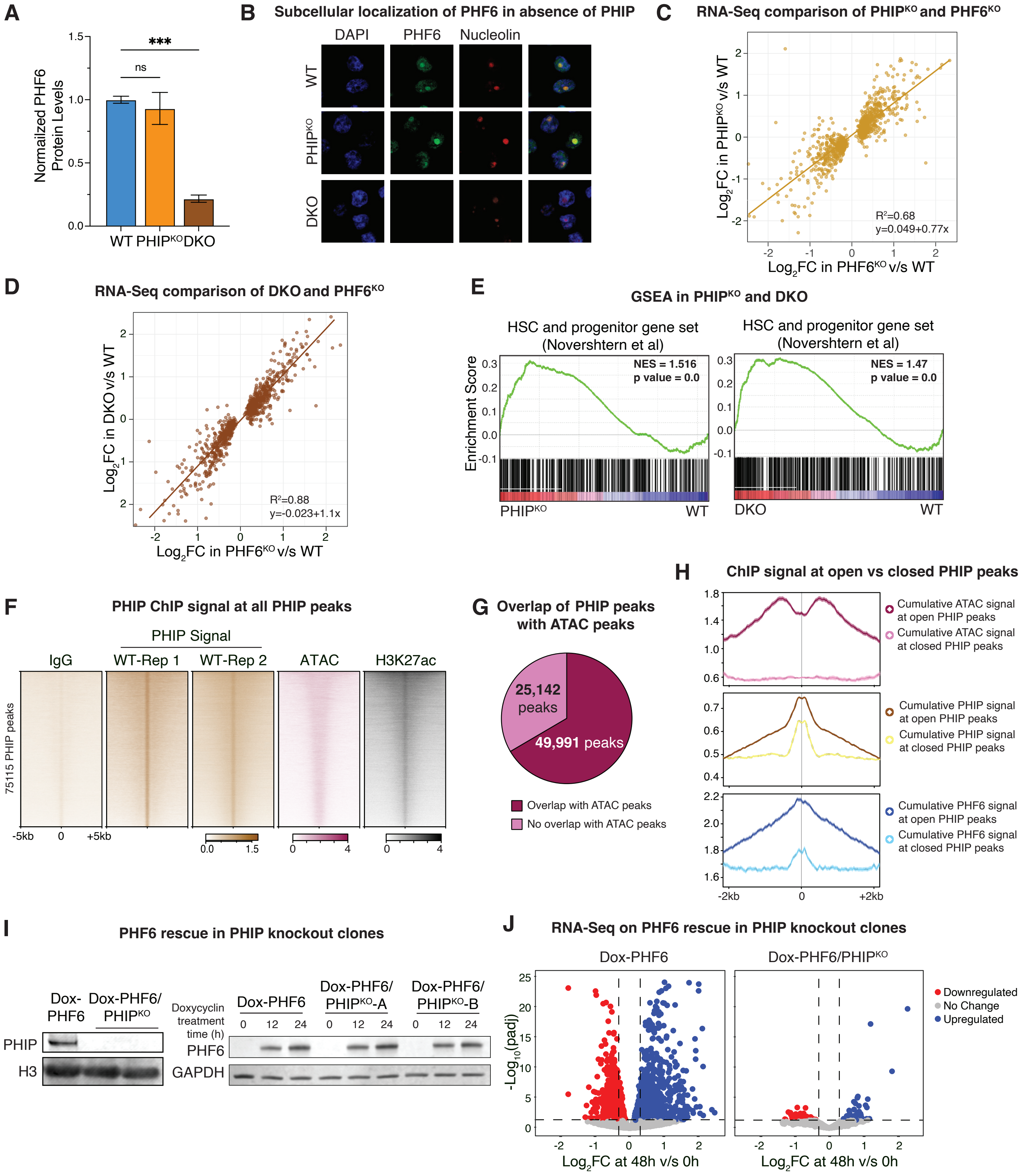

**Figure S5 (related to Fig 5)**

A. Bar graph showing PHF6 protein levels (normalized to GAPDH) in WT, $PHIP^{KO}$, and DKO clones. (n=3)

B. Representative immunofluorescence images showing localization of PHF6 protein in WT, $PHIP^{KO}$, and DKO clones. DNA stain DAPI marks the nucleoplasm, and nucleolin is a nucleolar marker.

C. Scatter plot showing positive correlation between effects of *PHF6* knockout and *PHIP* knockout on genes found to be differentially expressed in $PHF6^{KO}$ compared to WT.

D. Scatter plot showing positive correlation between effects of *PHF6* knockout and double knockout on genes found to be differentially expressed in $PHF6^{KO}$ compared to WT.

E. GSEA plot showing positive enrichment of HSC and progenitor cells gene set in $PHIP^{KO}$ (left) and DKO (right) compared to WT.

F. Heatmaps of two replicates of PHIP ChIP-Seq signal at PHIP peaks, along with ATAC-Seq and H3K27ac ChIP-Seq. IgG ChIP-Seq in WT is shown as negative control.

G. Pie chart showing distribution of PHIP peaks at open and closed regions of chromatin.

H. Metagene profiles of ATAC-Seq (top), PHIP ChIP-Seq (middle), and PHF6 ChIP-Seq (bottom) signal at open and closed PHIP peaks.

I. Immunoblot of PHIP in Dox-PHF6 and Dox-PHF6/$PHIP^{KO}$ clones (left). H3 is shown as loading control. Immunoblot showing time course rescue of PHF6 in Dox-PHF6 and Dox-PHF6/$PHIP^{KO}$ clones (right). GAPDH is shown as loading control.

J. Volcano plots showing differential expression of genes on PHF6 rescue in Dox-PHF6 and Dox-PHF6/$PHIP^{KO}$ clones. Dox-PHF6 volcano plot is the same as that shown in Fig S3C.

All bar graphs show mean ± standard error of mean (SEM), ns (not significant) = $p \geq 0.05$, *p = 0.01 to 0.05, **p = 0.001 to 0.01, ***p = 0.001 to 0.0001, ****p < 0.0001, by one-way ANOVA with Sidak's multiple comparison testing.
